# Unsigned temporal difference errors in cortical L5 dendrites during learning

**DOI:** 10.1101/2021.12.28.474360

**Authors:** Gwendolin Schoenfeld, Sepp Kollmorgen, Matthias C. Tsai, Christopher Lewis, Shuting Han, Philipp Bethge, Anna Maria Reuss, Adriano Aguzzi, Walter Senn, Valerio Mante, Fritjof Helmchen

**Affiliations:** Laboratory of Neural Circuit Dynamics, Brain Research Institute, University of Zurich, Zurich, Switzerland; Institute of Neuroinformatics, University of Zurich and ETH Zurich, Zurich, Switzerland; Institute of Neuropathology, University Hospital Zürich and University of Zurich, Zurich, Switzerland; Neuroscience Center Zürich, University of Zurich and ETH Zurich, Zurich, Switzerland; University Research Priority Program (URPP), Adaptive Brain Circuits in Development and Learning (AdaBD), University of Zurich, Zurich, Switzerland; Computational Neuroscience Group, Department of Physiology, University of Bern, Bern, Switzerland

## Abstract

Learning goal-directed behavior requires association of pertinent sensory stimuli with behaviorally relevant outcomes. In the mammalian neocortex, dendrites of pyramidal neurons are suitable association sites^1–3^ but how their activities adapt during learning remains elusive. Here, we track calcium signals in apical dendrites of layer 5 (L5) pyramidal neurons in mouse barrel cortex during texture discrimination learning^4^. We observe diverse task-related responses, either localized to branches or widespread throughout the apical tuft. However, even in expert mice, the tufts’ capability to discriminate go/no-go stimuli remains poor. Yet, we identify two prevailing response types in dendritic branches: 1) responses to unexpected outcome (reward) in naïve mice that decrease with growing task proficiency, and 2) responses associated with salient sensory stimuli, especially the outcome-predicting texture touch, that strengthen upon learning. We demonstrate that these response types match distinct unsigned components of the temporal difference error^5^ by replicating our results with a reinforcement learning model. Moreover, optogenetic apical inhibition of L5 neurons during the outcome time window prevents naïve animals from learning the task, consistent with the effect of equivalent model perturbation. Our findings suggest that salience signals in L5 apical dendrites facilitate the recruitment of task-relevant neurons via dendritic gain modulation.

## Introduction

Learning a new behavior requires the integration of diverse information—environmental context, sensory cues, specific stimuli, and associated outcomes (e.g. reward)—to drive suitable adaptations of brain activity and establish outcome predictions and appropriate motor actions. Accordingly, neuronal circuits can identify relevant stimuli and dynamically reorganize their activity upon task learning^2^. For example, neuronal populations in the whisker-related barrel field of primary somatosensory cortex (S1), especially those projecting to secondary somatosensory cortex (S2), undergo functional changes that reflect behavioral adaptations during texture discrimination learning^6^. Likely, these changes are implemented through plasticity mechanisms at the synaptic and cellular levels, including adaptations in neuronal dendrites^7^. The complexity of dendrites with their nonlinear properties makes them powerful computational elements and prime candidate sites for learning-related adjustments^3,8,9^. The distal apical dendrites of neocortical pyramidal neurons reach into layer 1 (L1), which receives rich and diverse inputs from local as well as distant sources^10–12^. Distal dendritic activity can reflect sensory stimuli^13,14^, motor actions^15,16^, feedback from higher-order thalamic^17,18^ or cortical^19,20^ areas, and reward^21^. Computationally, apical dendrites may impact somatic activity through several mechanisms, including gain modulation^22^, coincidence detection of somatic and distal dendritic activity^8^, and prediction error signalling^23–25^. Through such mechanisms they could help to solve the credit assignment problem^26^. Yet, a unifying concept of the computational role of apical dendrites in learning-related brain circuit adaptations is missing^22^. Here, based on experimental evidence and modeling results, we put forward the concept that L5 apical dendritic signals reflect distinct unsigned prediction error terms of temporal difference (TD) learning. We propose that apical dendrites convey unconditioned and conditioned salience information to the neurons and thereby promote the recruitment of task-relevant neurons via dendritic gain modulation.

## Results

### Chronic imaging of L5 apical tuft activity during learning

We sparsely expressed GCaMP6f in L5 pyramidal neurons in S1 barrel cortex of 6–10-week-old adult Rbp4-Cre mice. We specifically targeted S2-projecting L5 neurons by using an intersectional Cre/Flp viral labeling approach (Fig. 1a; Methods). Because Rbp4-Cre mice are not specific for L5 subtypes, labeled neurons presumably included L5A and L5B neurons sparsely distributed across barrel cortex (Supplementary Fig. 1). We trained mice (n = 8) in a go/no-go texture discrimination task^4,6^ (Fig. 1b; Methods). In each trial, we presented either a coarse (grit size P100) or a smooth (P1200) sandpaper to the whiskers (go-texture: P100, n = 3 mice; P1200, n = 5). Correct licking in go trials (Hits) triggered a water reward, whereas correct rejections (CR) in no-go trials (and Misses in go trials) were neither rewarded nor punished. False alarms (FA) in no-go trials were mildly punished with acoustic white noise. For analysis, we defined four time windows linked to the trial structure (auditory cue, touch, late-touch, and outcome; Fig. 1c; Methods). All mice improved their performance from naïve state (50% chance level) to expert level (>75%; Methods), on average requiring 1178 ± 368 trials (8-14 days; 1 session per day; 67-209 trials per session; mean ± s.d.; n = 8 mice). To compensate for different learning rates, we aligned all individual learning curves to the first expert trial (Fig. 1d) and focused our analysis on the training period from 500 trials before to 150 trials after this time point (trial identifier [ID] from −500 to 150). Based on identified learning onset and first expert trial we divided the training period into ‘naïve’, ‘learning’ and ‘expert’ phase (Methods). With learning, mice developed anticipatory whisking preceding the first whisker-texture touch as well as anticipatory whisking and licking before the outcome window^6^ (Supplementary Fig. 2a-c).

**Fig. 1.**
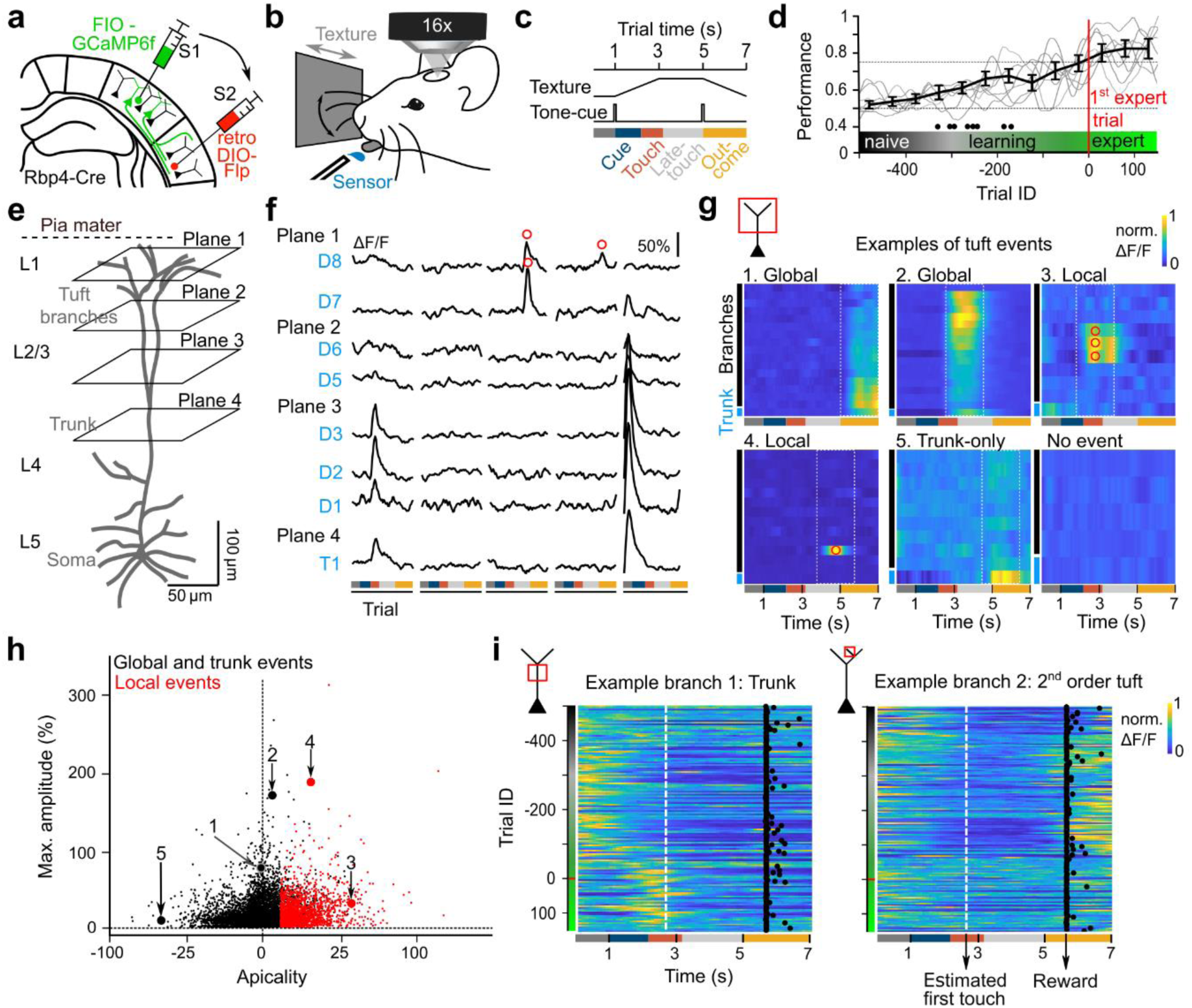
Longitudinal multi-plane calcium imaging of L5 dendrites across learning. **a**, Dual-virus approach in Rbp4-cre mice to express GCaMP6f in S1→S2 projecting L5 pyramidal neurons. **b**, Schematic of go/no-go texture discrimination task. **c,** Trial structure with defined time windows for analysis. **d**, Learning curves across 650 trials aligned to first expert trial (n = 8 mice). Black line: mean ± s.d. in 50-trial bins; grey lines: individual mice; black dots indicate transitions from naïve to learning period for individual mice **e**, Schematic of multi-plane two-photon imaging from 4 depths. **f**, Example calcium transients across all imaging planes in selected ROIs of trunk (T) and dendritic branches (D) for a single L5 tuft. Five example trials are shown, red circles indicate local events in distal branches. **g**, Heatmaps of tuft activation patterns (normalized ΔF/F) showing examples from the response spectrum (1: Trunk-dominated global event, 2: Tuft-dominated global event, 3: Local event in 3 branches, 4: Single-branch local event, 5: Trunk-only event, 6: Trial without event). Examples are from different tufts and each heatmap corresponds to one trial. Note the occurrence of events in different trial windows (vertical lines demarcate 2-s windows used for event detection). Red circles indicate local events. **h**, Distribution of tuft activation patterns. Each data point represents a 2-s event, for which a ΔF/F transient peak was detected. For each event, the maximal ΔF/F amplitude (min-to-max range) across tuft branches is plotted versus the ‘apicality’ of the event, defined as the binary logarithm of the tuft/trunk amplitude ratio (Methods; local events required apicality > 2). Numbers and arrows mark the events shown in panel g. **i**, Heatmaps of normalized ΔF/F traces across learning for two example dendrites (a trunk and a 2^nd^ order tuft branch). Dashed vertical line indicates approximate time of first touch. Black dots indicate reward valve opening times. Note that the trunk exhibited early activity during trials that shifted closer to the texture stimulation time during learning, whereas the tuft branch displayed late activity in the outcome window that diminished upon learning.

To investigate dendritic activity during learning, we applied multi-plane two-photon calcium imaging and repeatedly measured calcium transients (ΔF/F) in dendritic tuft branches and parent apical trunks of S1→S2 L5 neurons throughout the training period (Fig. 1e; Supplementary Fig. 3; 30-370 µm depth below the pia; effective frame rate 10 Hz; Methods). We manually selected trunk cross-sections and dendritic branch segments (continuous stretches of 5-20 µm) as regions of interests (ROIs). Multi-plane calcium imaging enabled us to simultaneously record activity in multiple compartments of the same apical tuft (Fig. 1f). We observed diverse tuft activation patterns, including ‘global’ events, i.e., concurrent large-amplitude signals in the trunk and several or all tuft branches, and ‘local’ events, i.e., clearly detectable calcium signals in only one or few tuft branches with no detectable signal in the trunk (Fig. 1g). In addition, large trunk signals could occur without clear activation of tuft branches (‘trunk-only event’). Global events likely are associated with dendritic calcium spikes and somatic action potential bursts^27,28^, whereas local events in tuft branches may reflect distinct forms of dendritic spikes^20,29^ that were not sufficiently strong, or lacked coincident somatic excitation, to trigger calcium spikes near the main bifurcation.

For quantification, we detected tuft events based on local peaks in a 2-s sliding window across all trials and categorized them into local events versus global or trunk events (Fig. 1h; see Methods for criteria). Individual apical tufts could show local events with distinct activity patterns, i.e., engaging different subsets of tuft branches, which sometimes re-occurred in multiple trials across days (Supplementary Fig. 4). Overall, we detected at least one tuft event in 28.5% of all 24’330 recorded trials, out of which 24.2% were classified as local events (1678 of 6943 detected events). Thus, local tuft events were relatively frequent in S1 L5 neurons under our conditionsbut with high variability. In view of this high variability, we decided to focus our analysis of learning-related changes mainly on the single-branch level, without considering whether an individual branch signal related to a local event or occurred as part of a global event. As exemplified by ΔF/F traces from the trunk and a tuft dendrite belonging to the same L5 example neuron (Fig. 1i), dendritic branches differed not only in the timing when they were preferentially active during trials, but also in the dynamic changes of their activity profiles across learning. These learning-related changes in dendritic activity did not correlate with the observed behavioral changes in whisking and licking (Supplementary Fig. 2d,e), suggesting a more specific role in learning dynamics.

### Two major types of learning-associated dendritic response dynamics

For a comprehensive analysis of task- and learning-related dendritic dynamics in our entire data set, we identified and selected trials with significant ΔF/F traces (Methods). For visualization we used a UMAP embedding based on the similarity of the trial-related ΔF/F traces. Both the short timescale (within single trials) and the long timescale (across trials) are reflected in the UMAP plot (Fig. 2a): First, color-coding according to the onset times of clearly detectable dendritic calcium transients (Methods) shows that dendritic activity is distributed over the entire trial time course, including cue, touch, late-touch, and outcome windows (upper plot, Fig. 2a). A small fraction of traces displayed transients with double peaks in both cue and outcome windows, represented in the middle of the UMAP plot. Second, color-coding based on trial ID revealed an interesting pattern of overrepresentation of calcium signal onsets around the touch window start in expert trials, whereas in early naïve trials calcium transients in the cue and outcome windows appeared more abundant (lower plot, Fig. 2a; for color-coding of other features see Supplementary Fig. 5). This overview indicates that apical dendritic activity in S1→S2 L5 neurons reflects various task-related events and that the trial-related temporal pattern of dendritic activation reorganizes during learning.

**Fig. 2.**
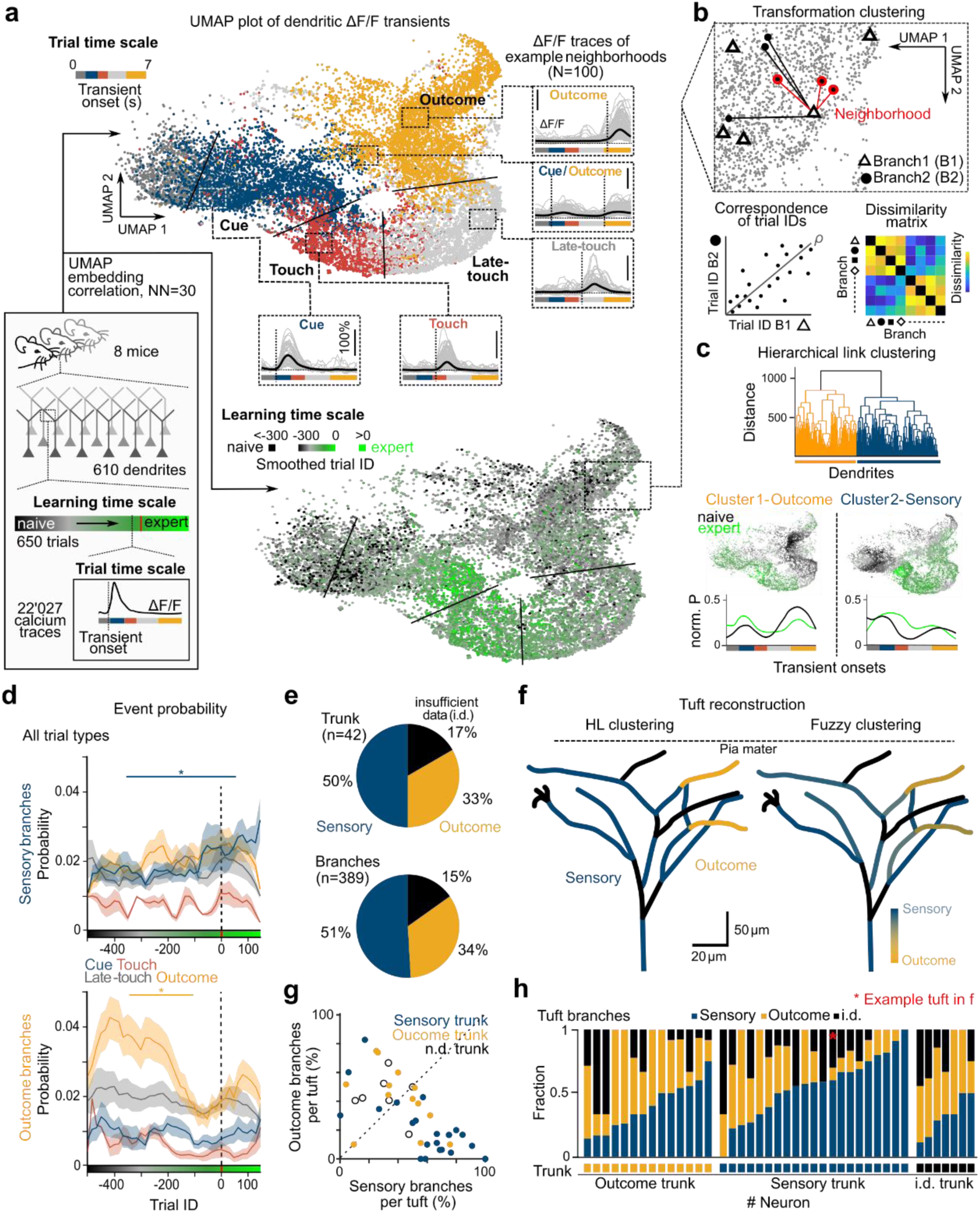
Two dendritic subclasses defined by distinct changes in sensory and reward representations. **a***, Top:* Correlation-metric-based UMAP embedding of all recorded ΔF/F traces with detectable transient onsets, colored according to transient onset time. Inserts: ΔF/F traces and average ΔF/F traces associated with local 2D neighborhoods. *Bottom:* UMAP embedding colored according to smoothed trial ID (8 mice, 22’027 traces). **b**, *Top*: Schematic depiction of the transformation clustering method. **c**, Hierarchical link clustering identifying of two functionally distinct clusters of dendritic responses by their change in ΔF/F transient onset distribution over learning. *Middle:* UMAP embedding of data points belonging to the respective cluster colored by smoothed trial ID (see Fig. 2a bottom). Bottom: Smoothed normalized distribution of calcium transient onset times across trial time in the naïve and expert conditions for both clusters. **d**, Transient onset probability during learning in cue, touch, late-touch and outcome window in sensory-driven and outcome-driven class (Sensory branches: Cue window: naïve vs. expert: p= 0.03; Late-touch: naïve vs. expert: p=0.05. Outcome branches: Outcome window: naïve vs. learning: p=0.02.). **e**, Percentages of trunks and apical branches per class. Branches with insufficient data (i.d.) were not assigned to a class. **f**, Reconstruction of example tuft from a two-photon image stack. *Left:* Branches are colored according to the respective HL class. *Right:* Branches are colored according to fuzzy c-means clustering. **g**, Class assignment of apical trunks in relationship to the percentage of branches classified as sensory and outcome. **h**, Fraction of branches per neuron assigned to each cluster (42 tufts; sorted according to trunk class, sub-sorted according to percentage of sensory branches).

To evaluate this functional reorganization of dendritic dynamics quantitatively and to examine whether consistent learning-related patterns exist, we developed ‘transformation clustering’, a statistical approach to identify dendritic branches (including apical trunks) that exhibited similar learning-associated activity changes. Specifically, we compared the high-dimensional single-trial responses (not the dimensionality-reduced UMAP data points) for all possible dendrite pairs across learning using a nearest-neighbour method^30^ and analyzed the resulting dissimilarity matrix with hierarchical link (HL) clustering (Fig. 2b; we greedily expanded clusters by also assigning dendritic ROIs with sparse activity; Methods and Supplementary Fig. 6). We identified two major functionally distinct types of dendritic activity changes across learning. One type (Cluster 1; referred to as ‘outcome’ type) shows abundant and large calcium transients in the outcome window for rewarded Hit trials in naïve mice and a reduced probability of these outcome-related responses in experts (Fig. 2c,d). These changes are evident in the average onset distributions of ΔF/F transients across trial time as well as across learning (Fig. 2c; they are less obvious in averaged ΔF/F traces, though; Supplementary Fig. 7). The second functional type (Cluster 2 or ‘sensory’ type) comprises dendritic branches that display an increasing probability in expert mice of ΔF/F transients during the cue and touch windows (Fig. 2c,d). This response type apparently highlights time windows of salient task events, particularly the reward-predicting texture touch. The distinct temporal dynamics of outcome and sensory response types across learning are also evident in the respective UMAP subplots (Fig. 2c and Supplementary Fig. 5).

To analyze the tuft composition in terms of functional types, we evaluated 42 neurons with identified trunk ROIs. For both trunks and tuft dendrites, about half belonged to the sensory class and about one third to the outcome class as determined by HL clustering (Fig. 2e). The remaining dendrites could not be assigned to a class because of insufficient number of data points (i.d.). Additional to the HL clustering we performed fuzzy c-means clustering, revealing that the two dendritic classes represent two extremes of a continuum of dendritic activity patterns (Fig. 2f; Supplementary Fig. 8). In 38 of 42 neurons the tuft contained dendritic branches of both functional types, with the functional type of the trunk largely matching with the most abundant type in its tuft branches (Fig. 2g,h; see also ΔF/F traces of example neurons in Supplementary Fig. 9). No difference in morphology was obvious in tufts dominated by either of the functional classes (Supplementary Fig. 10). Taken together, we identified two major functional types of dendritic responses with distinct changes in their activity profiles across learning. We interpret these response types as reflecting changes in the effective strengths of distinct input streams converging on individual L5 dendritic tufts, carrying information about context (auditory cue), relevant sensory stimulus (touch and late-touch), and reward (outcome window). We next asked what computational role these dendritic response types might have.

### Dendritic response types reflect distinct unsigned TD prediction error components

We first examined whether dendritic branches of the two types or entire tufts attain the ability to discriminate different trial types as reported for L2/3 neuronal somata^4,6,31^. To assess discrimination power, we trained a linear decoder on either whole-tuft or individual-branch data and evaluated its performance using receiver operating characteristic (ROC) analysis (Methods). Even when considering whole-tuft activity, no significant discrimination power emerged upon learning, neither for sensory-dominated tufts in the touch window nor for outcome-dominated tufts in the outcome window (Fig. 3a; Supplementary Fig. 11 for individual branch analysis). This result suggests that dendritic tufts may be less important for fine discrimination of the relevant stimulus but rather for accentuating time windows of salient task-related events: the unconditioned reward (or punishment) in the outcome window in the naïve state when outcomes are unexpected and the conditioned sensory stimuli (predictive auditory cue and whisker touch) when the animal has learned the task structure.

**Fig 3.**
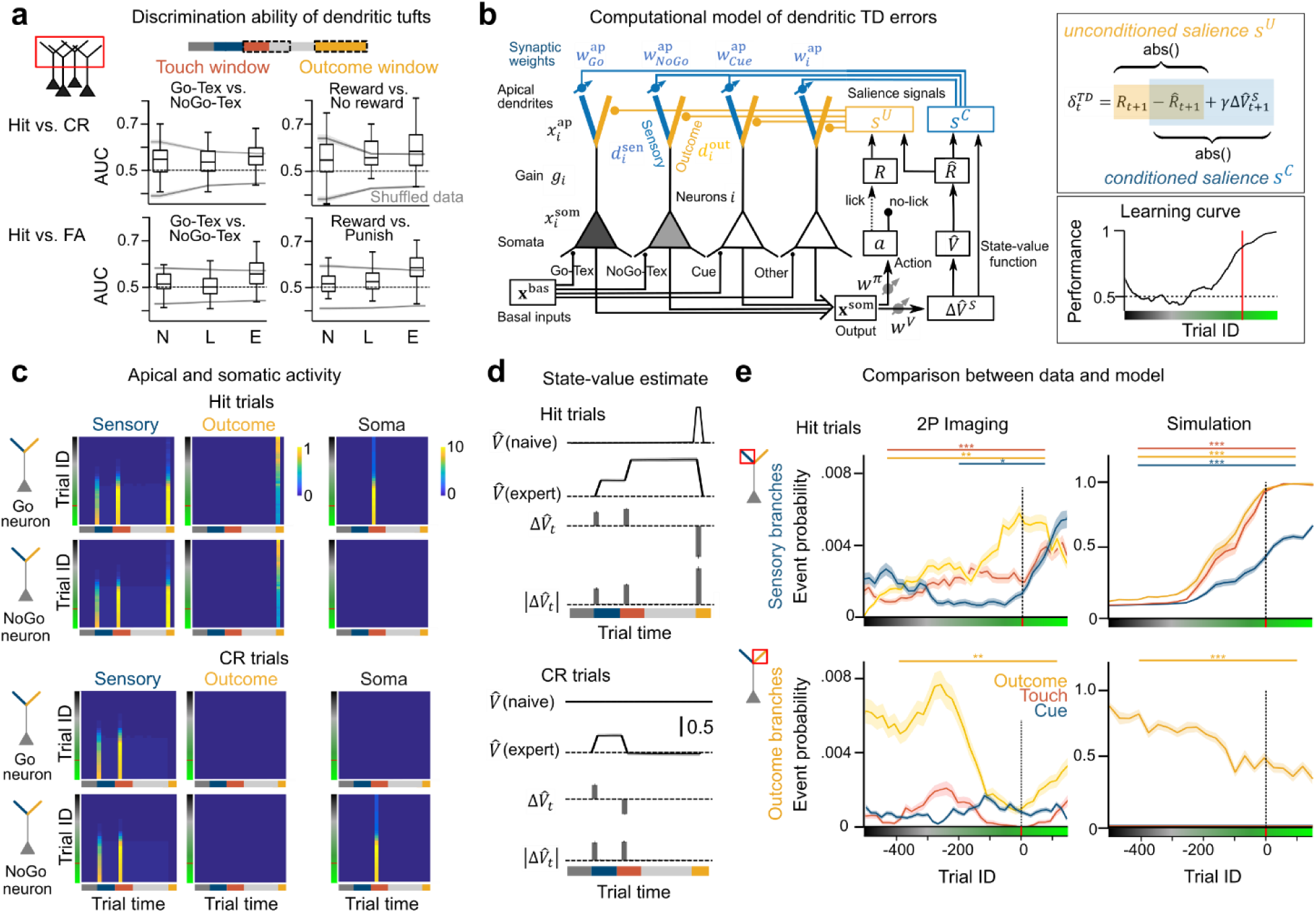
Computational model of salience learning predicts dendritic activity changes *in vivo*. **A,** Discrimination analysis of tuft ΔF/F patterns to discriminate sandpaper texture in the texture window (2.4-3.9 s) or trial outcome in the outcome window (5-7s) using a linear support vector machine (SVM) across all tufts in the naïve (N), learning (L), and expert (E) state. Grey lines indicate 95% and 5% quantile of shuffled data (mean ± s.e.m.). **b**, Left: Schematic of the computational model for salience-based TD-learning in S1 L5 neurons. See main text and Methods for details. Upper right: definition of the two unsigned components of the TD error, representing unconditioned and conditioned salience. For completeness, the parameter *γ* for discounting is included in the equation but we did not consider discounting (*γ* = 1). Lower right: example learning curve of the model. **c**, Simulation of activity across trial time and trial ID in the three compartments (sensory branch, 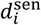; outcome branch, 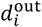; and soma, 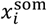) of Go and NoGo neurons during Hit trials. **d**, Depiction of the modelled state-value estimator *V̂*, its change over trial time and the absolute value thereof, corresponding to conditioned salience, for Hit (top) and CR (bottom) trials. **e,** Comparison between 2P imaging results (left) and model simulations (right) of ΔF/F calcium transient event probability in Hit trials per trial window across learning for sensory (top) and outcome (bottom) dendritic branches. Imaging data: 42 tufts, mean ± s.e.m; Sensory branches: Outcome window, naïve vs. expert, p=0.006; Touch window: naïve vs. expert, p<0.001; Cue window, learning vs. expert, p=0.02; Outcome dendrites: Outcome window, naïve vs. expert, p=0.004. For all significant comparisons in the simulations, p<0.001. *p<0.05, **p<0.01, ***p<0.001

We recognized that the two types of branch responses resemble two unsigned (salience) components of the classic temporal difference (TD) error that is used to update a state-value estimation^5^. Specifically, these components represent the unconditioned salience, *s*^*U*^, of unexpected outcome and the conditioned salience, *s*^*C*^, of task-relevant, outcome-predictive stimuli (Fig. 3b; Methods and Supplementary Note 1). We cast this idea into a computational model of a population of multi-compartmental L5 pyramidal neurons using TD reinforcement learning to train a state-value estimator function *V̂* (Fig. 3b; Supplementary Note 2). For each neuron *i*, 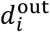 and 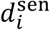 are the activities in two branches of the apical dendritic compartment, representing the two functional branch types (outcome and sensory). The sum of **d**^out^ and **d**^sen^ determines the apical dendritic activities **X**^ap^, which modulate the multiplicative gains **g** that govern the somatic activities **X**^som^ (Fig. 3b; Supplementary Fig. 12b-c). **d**^out^ is driven uniformly across all dendritic tufts with a fixed input strength by an input representing the unconditioned salience, defined as the absolute value of the reward prediction error, *s*^*U*^ = |*R* −*R̂*|. The conditioned salience input is learned to represent the unsigned temporal changes in *V̂*, *s*^*C*^ = |Δ*V̂*| = |−*R̂* + *γ*Δ*V̂*^*S*^| across trial time, which comprises changes due to expected outcomes when time progresses (−*R̂*) and relevant sensory stimuli as part of state changes (Δ*V̂*^*S*^) (Fig. 3b; Supplementary Note 1; *γ* is a discounting parameter). *s*^*C*^ is imposed on the sensory branches to drive 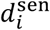 via neuron-specific plastic apical weights 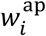. The plasticity of these synapses is governed by local coincidence detection between postsynaptic activity and dendritic inputs such that task-relevant neurons experience synaptic potentiation and task-irrelevant synaptic depression (Supplementary Fig. 12, Supplementary Note 3). The state-value function estimator *V̂* and the action policy (lick or no-lick) are learned by adjusting synaptic weights *w*^*V*^ and *w*^*π*^ that connect the somatic output of the L5 population to both a Δ*V̂*^*S*^ estimation network and an action selection network (Fig. 3b; Supplementary Note 2). We presume that the state-value estimation and action selection occur in neural circuits outside of barrel cortex that establish value representations and motor plans during learning, likely involving frontal cortical regions^32^.

In the model, the feedforward basal inputs **X**^bas^ onto L5 somata are fixed throughout training and do not need adjustment. For simplicity, we first assumed one go-texture (Go) neuron and one no-go-texture (NoGo) neuron, which receive basal input in the touch window exclusively in go and no-go trials, respectively (pure-selectivity model). In addition, we presumed one tone-cue (Cue) neuron that receives basal input in the tone-cue window in all trials, and 200 distractor neurons that receive noisy random basal inputs (Supplementary Figure 12a; Methods and Supplementary Note 2). As the salience signals in the apical tufts develop, the somatic response of the Go neuron is amplified in Hit trials through dendritic gain modulation and associated to the lick action. Correspondingly, the NoGo neuron is amplified in CR trials and associated with not licking (Fig. 3c; Supplementary Fig. 13). Note that both 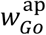 and 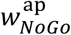, as well as 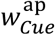, increase during learning, because both texture stimuli and the tone-cue are relevant for the task and affect outcome prediction. In contrast, the apical weights of the task-irrelevant distractor neurons are reduced, lowering their somatic activities. As a hallmark of TD-learning and conditioning, the initial dendritic response to a reward signal is advanced in time so that after learning the response is already elicited by the presentation of the conditioned texture stimulus, and even earlier at the time of the conditioned auditory tone-cue (Fig. 3c,d). The conditioned salience signal *s*^*C*^ = |Δ*V̂*|, driving the sensory dendrites of task-relevant neurons in the expert state, are interpreted as temporally advanced reward-prediction errors that highlight moments, at which relevant stimuli (texture, tone) are preceived and thus lead to an update of the state-value estimation *V̂* (Fig. 3d). Comparing the model dynamics across learning with our experimental findings, we find good agreement of the learning-related modulation of sensory and outcome dendrites (Fig. 3e).

In addition to the pure-selectivity model, we also tested a more general mixed-selectivity model, in which basal inputs **X**^bas^ provide mixed sensory inputs to the L5 somata, resulting in a distribution of go/no-go texture preference across the L5 population. This model also learned the task by amplifying the activity of the most task-relevant neurons, i.e., those that received inputs biased to either go or no-go stimulus. As a result, their somatic texture selectivity was enhanced (Extended Fig. 14). Overall, we put forward a new framework for the computational role of apical tuft dendrites in learning, emphasizing the potential importance of adjusting dendritic gain modulation across the population such that at each salient time point the appropriate neurons are recruited.

### Inhibition of outcome-window activity impairs learning

We hypothesized that the large signal in the outcome window in naïve mice (the dominant TD error component at this stage) might be essential for driving circuit adaptations and thus for learning. To assess whether the unconditioned salience signal conveyed via dendrites of S1→S2 L5 neurons is relevant for learning, we densely expressed eArchT3.0 in S1→S2 L5 neurons to optogenetically suppress apical dendritic activity during the outcome window (Fig. 4a). We validated the inhibitory effect of laser stimulation on eArchT3.0-expressing L5 neurons using extracellular recordings in anesthetized mice: Direct surface illumination reduced but did not eliminate spontaneous multi-unit activity and generated a pronounced sink in the current source density (CSD) signal in superficial layers including L1 (Fig. 4b; Supplementary Fig. 15a). The effect of surface stimulation was comparable to superficial illumination targeted to L1 through an oblique optical fiber, whereas illumination targeted to deep layers nearly abolished both spontaneous and whisker-evoked spiking activity in L5 (Fig. 4c,d; Supplementary Fig. 15b,c).

**Fig. 4.**
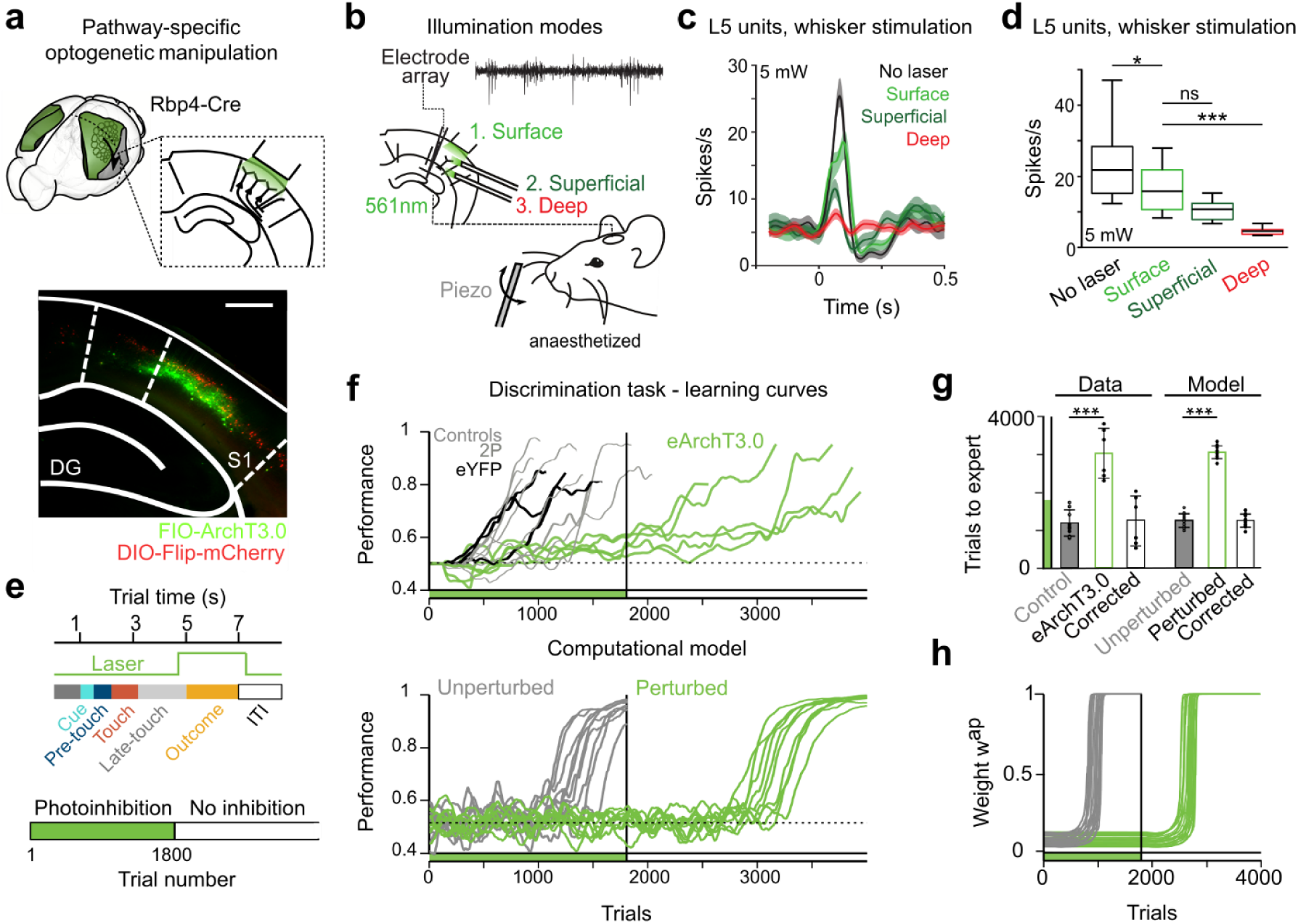
Optogenetic suppression of outcome-driven activity prevents learning. **A,** Pathway-specific optogenetic manipulation of S1→S2 L5 neurons using eArchT3.0. *Bottom:* Dense labeling of eArchT3.0 in L5 neurons (confocal image, coronal view, 100 µm). **b**, Schematic of multiple compartment optogenetic perturbation using three fiber optic cannulae with simultaneous electrode shank recording in isoflurane-anesthetized mice during whisker stimulation. Three different types of illumination were tested: surfaces illumination (as in the awake experiment), superficial illumination aimed a L1 and deep illumination at L5. **c**, Whisker stimulation evoked firing rate response of an example L5 unit that showed suppression for perturbation of different compartments (mean ± s.e.m.). **d**, Firing rate of L5 units with and without optogenetic manipulation according to fiber position (line: median, box: 25th and 75th percentile, whiskers: 5th and 95th percentile; 5 mW, 17 units, 200 trials, 8 sessions, n = 4 mice, t-test *p<0.0025, ***p<0.001). **e**, Temporal profile of optogenetic inhibition during trials (top) and across training (bottom). **f**, Top: Optogenetic perturbation during a texture-discrimination task was carried out for 1’800 trials. Learning curves of eArchT3.0-expressing mice (n = 5) and control mice (from GCaMP6f experiments, n = 8; and eYFP controls, n = 3)). Bottom: Computational model simulation of learning curves for unperturbed (grey, n = 10) and perturbed (green, n = 10) agents. For the perturbed agents, the apical dendritic activity was set to zero and the adaptation during the outcome window blocked for 1’800 trials **g**, Left: Average number of trials required to reach expert performance in unperturbed mice (Control), optogenetically perturbed eArchT3.0 mice (eArchT3.0), and for perturbed mice after subtraction of the 1’800 perturbed trials (Corrected; 1’150 ± 314, 3’019 ± 659, and 1’219 ± 659 trials, respectively; mean ± s.d.; p<0.001 for Perturbed vs. Control; p = 0.89 for Corrected vs. Control; Wilcoxon rank-sum test). Right: Average number of trials required to reach expert performance of simulated agents in the model under unperturbed conditions (Unperturbed), with perturbation (Perturbed), and for the perturbed agents after subtraction of the 1’800 perturbed trials (Corrected) (1’239 ± 176, 3’043 ± 168 and 1’243 ± 168 trials, respectively; mean ± s.d.; p<0.001 for Perturbed vs. Unperturbed; p = 0.88 for Corrected vs. Unperturbed; Wilcoxon rank-sum test).**h**, Synaptic strengthening of apical synapses mediating the conditioned saliency input in perturbed and unperturbed agents (mean ± s.e.m.; n=10).

These results indicate that at intermediate laser intensity (5 mW; Supplementary Fig. 15d,e) optical stimulation induced mainly apical inhibition without blocking somatic spikes, comparable to similar previous approaches^13,16^.

Having verified the perturbation effect on L5 neuronal populations in the anesthetized condition, we applied 5-mW optogenetic inhibition during the 2-s outcome window in awake mice (n = 5) in every trial, starting in the naïve condition and continuing during texture discrimination training (Fig. 4e). We perpetuated this photoinhibition for 1’800 trials, which is about twice the average number of trials that mice normally required to reach expert performance. At the end of this long period of photoinhibition, none of the eArchT3.0-expressing mice had reached expert level (mean performance 62% ± 6.2%). After 1’800 trials, the optogenetic block was lifted, and only then eArchT3.0-expressing mice were able to improve their task performance to expert levels, with a time course comparable to mice expressing eYFP or GCaMP6f (Fig. 4f,g; Supplementary Fig. 15e). Licking and whisking behavior remained unaffected by the optogenetic perturbation (Supplementary Fig. 15f-i). These findings are consistent with our TD learning model, where blocking dendritic activity (**X**^ap^ = 0) in the outcome window for the first 1’800 trials prevented learning (Fig. 4f,g). Only after terminating this blockade, the trained agents were able to learn the task and to reach expert performance in a comparable number of trials as the optogenetically perturbed mice. The reason is that only after the end of the blockade, synaptic weights 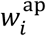 conveying the conditioned salience signal were plastically adjusted (Fig. 4h) so that the appropriate subset of sensory branch signals 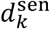 could be strengthened and exert their modulatory effect on somatic activity. Taken together, our experimental and modeling results suggest that the unconditioned salience information inducing apical dendritic activity of S1→S2 neurons in the outcome window is essential for learning the new task.

## Discussion

Here we discuss the role of apical dendrites in shaping neural activity patterns during learning, based on the results of our longitudinal study of dendritic tuft calcium signals in L5 pyramidal neurons. We link these results to a TD learning model that utilizes unsigned prediction errors, and we propose a general concept, in which the activities of task-relevant neurons are amplified through learned dendritic gain modulation. Such amplification may be instrumental for establishing and promoting appropriate signal flow through large-scale neural circuitry, tailored to highlight and make use of the information most salient to the animal.

Besides widespread tuft activation events, we observed dendritic events localized to one or few branches, similar to several previous studies^15,20,33,34^, but contrasting other studies^35,36^ that reported highly correlated activity in soma and apical dendrites of L5 neurons. The reasons for these discrepant results are unclear, possibly reflecting differences in cortical region, neuronal subtype, behavioral state, or task demands. The prevalence and relevance of local dendritic branch activity in L5 tufts *in vivo* thus warrants further investigation^37^.

Despite the variability of dendritic signals, we identified two functional types of branch activity, distinguished by their timing during task trials and their distinct profiles of learning-related adaptation. We interpret these response types as reflecting converging input streams that convey two distinct types of salience information to the apical tufts. Individual branches are dominated by one or the other input type, possibly reflecting local connectivity including potential clustering of synaptic contacts^38^. We formalized the notion of salience processing and replicated our results in a TD learning model, in which unsigned TD error components reflecting unconditioned and conditioned salience are fed onto dendritic tuft branches (outcome and sensory branches, respectively). Dendritic responses to both unexpected random rewards and trial-associated rewards have been previously reported in L5 tufts of barrel cortex^21^. Reward prediction errors carrying unconditioned salience information may originate from various sources, including prefrontal areas linked to the midbrain dopaminergic system, as well as direct cholinergic^39,40^, serotonergic^41^ or noradrenergic^42,43^ modulatory afferents. Potential sources of conditioned salience information, on the other hand, include perirhinal cortex conveying stimulus-outcome associations^19,39^, higher-order thalamic afferents^44–47^, posterior association areas conveying contextual or anticipatory information ^19,48,49^, and inputs from frontal premotor and motor areas^14,50,51^. Furthermore, areas of the salience network^52^ could be involved, such as anterior cingulate cortex, agranular insular cortex and mediodorsal (MD) thalamus, which all display projections to L1 in barrel cortex^53^.

All these projections from various regions might affect L5 apical tuft dendrites directly or indirectly by synapsing onto local inhibitory interneurons. Notably, widespread reward-associated activation of vasoactive intestinal polypeptide(VIP)-expressing interneurons^54^ leads to disinhibition of L5 apical dendrites, likely opening windows of opportunities for other pathways to modulate dendritic activity in the outcome period. Likewise, stimulus-associated activation of L1 interneurons might be essential in gating and regulating the impact of long-range inputs on apical dendrite activation^47,55,56^. Generally, the learning-related occurrence of unconditioned responses before the conditioned response appears to happen in a widely distributed manner^57^. In S1, dendritic response adaptations might involve local plasticity at L1 synapses (in tuft dendrites themselves or in surrounding interneurons) or plasticity in further upstream regions such as S2^31^, perirhinal^19^, premotor^51^, or orbitofrontal^58^ cortices and other regions that might be involved in the formation of state-value representations. Fully dissecting the local and long-range circuitry involved in the learning-related adaptations of L5 neuronal function remains a paramount objective.

The theoretical framework derived from our experimental data brings together three computational facets that have been associated with dendritic function for decades: gain modulation, prediction error signaling, and somato-dendritic conincidence detection. Gain modulation of neurons^56^, particularly via dendritic activity^22^, is a powerful means to dynamically adjust a neuron’s output in the light of changing input. Here, we propose multiplicative gain control^56^ as a mechanism to amplify specific neuronal subsets that, in a given context, encode valuable information to solve a goal-directed task. The mechanism by which neurons are selected combines the other two facets. The activation of distal apical dendrites of task-relevant neurons by learned salience signals that relate to TD error components opens time windows of opportunities, during which somatic activity can be amplified. Interestingly, the task relevance of a neuron *i* turns out to be appropriately defined by the covariance of *s*^*C*^ and 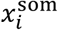 (see Supplementary Note 3), i.e., the covariance between the top-down salience signals conveyed via the apical dendrite and the somatic activity, highlighting the importance of dendro-somatic coupling and linking our idea to the notion of coincidence detection in apical dendrites^8,59,60^. The general concept behind this definition is: the more a neuron’s activity co-varies with the salience signals broadcasted via the apical tufts, the more likely it is to carry task-relevant information that can contribute to outcome prediction. Consequently, in the given context and task setting, such neurons with high apparent relevance should be engaged by dialing up their activity. Our photoinhibition experiments indicate that learning abilities are compromised when these mechanisms are suppressed in a relevant brain region.

In summary, we discovered that task- and learning-related dynamics of L5 apical dendrites reflects unsigned TD error components. We started mapping mathematical terms of a learning theory to specific neuronal processes, which is a promising avenue to be followed in future studies. Such mapping may not only reveal principles of neural dynamics underlying learning but may also inspire new designs of artificial neural networks^61^.

## Acknowledgements

This work is supported by grants from the Swiss National Science Foundation (project grants 170269 and 192617 to F.H., Sinergia grant 180316 to F.H. and W.S., grant 179040 to A.A, Award PP00P3-157529 to V.M., Ambizione grant 216312 to S.H.), the European Research Council (ERC Advanced Grant BRAINCOMPATH, project 670757, to F.H.), the University Research Priority Program (URPP) ‘Adaptive Brain Circuits in Development and Learning’ (AdaBD) of the University of Zurich (to F.H. and V.M.), a Forschungskredit from the University of Zurich (project K-41220-04, C.L.), NOMIS Distinguished Scientist Award to A.A., the Simons Foundation (SCGB 328189 and 543013 to V.M.), and the Horizon 2020 European Framework Programme under grant agreements 785907 and 945539 (HBP) for W.S. The authors thank Henry Lütcke for his support with the CaImAn pipeline, Ladan Egolf for managing transgenic mouse lines, Dubravka Dujmovic-Göckeritz for genotyping and CLARITY clearing of whole brains, and Nikita Vladimirov for help with mesoSPIM light-sheet imaging.

## Author contributions

G.S. and F.H. conceived the project and designed the study; G.S. carried out all awake *in vivo* experiments; C.L. performed extracellular recordings in anesthetized mice; G.S., S.K., and C.L. analyzed data; S.K. developed transformation clustering; A.M.R. performed iDISCO clearing of brain hemispheres; P.B. and A.M.R. performed light-sheet microscopy on cleared brains; M.T., F.H., and W.S. developed the computational model; M.T. implemented the model simulations; A.A., W.S., V.M., and F.H. supervised experiments and analysis; G.S., M.T. and F.H. wrote the manuscript with comments from all authors.

## Supplementary Information

is provided below (Supplementary Figures 1-15) and in 2 additional separate files (Supplementary Video 1 and Supplementary Notes 1-3).

## Methods

All experimental procedures were carried out in accordance with the guidelines of the Federal Veterinary Office of Switzerland and were approved by the Cantonal Veterinary Office in Zurich under license number 234/2018.

### Animals and preparations for chronic imaging

We used male and female adult 6-10 week old Rbp4-Cre transgenic mice (n = 8, Tg(Rbp4-cre)KL100Gsat/Mmucd, MGI:4367068, ref. ^62,63^). For surgical preparation, mice were anesthetized using isoflurane (1.5-2% in O_2_) and the body temperature was maintained at 37°C using a heating pad with rectal probe. After exposing the skull, a 4-mm diameter craniotomy was made above the left S1 barrel cortex and S2. Stereotactic injections of AAVretro-hSyn1-chI-FLEX-mCherry_2A_NLS_FLPo virus solution (6.3 × 10^12^ vg/ml, dilution 1:50) was injected into S2 (three injections à 210 nl; AP|ML|DV coordinates from bregma (in mm): −0.7|3.5|-1, −1.2|4.1|-1.5, −1.3|4.5|-1.5). AAV-2.1-hSyn1-fio-GCaMP6f virus solution (1.8 × 10^12^ vg/ml) was injected into L5 of barrel cortex (three injections à 210 nl: −0.7|-3|-0.6, −1.1|-3|-0.5, −1.1|-2.4|-0.6). The craniotomy was sealed with a 4-mm glass cover slip and dental cement (Tetric EvoFlow). A light-weight head-post was fixed on the skull using dental cement. For the 3 days following the surgery, animals were monitored and analgesics (Metacam, 5 mg/kg, s.c.) and antibiotics (Baytril, 10 mg/kg, s.c.) were administered. Animal handling began 5 days after surgery and the first imaging session took place >21 days after virus injection.

### Behavioral task and mouse training

The setup for the go/no-go texture discrimination task has been described previously^4,64^. Each trial started with the opening of the laser shutter (Thorlabs, SH05/M) followed after 1s by an auditory tone (two 2-kHz beeps of 100-ms duration with 50-ms interval). Then, either the rough or smooth texture (P100/P1200 sandpapers) was moved towards the whiskers on the right side of the animal’s snout. We presented the two texture types randomly but with no more than 3 repetitions. In expert mice, which typically show anticipatory whisking, the first texture-whisker touch typically occurs around 0.5 s before the texture stops^64^. After a 2-s stimulus presentation period the texture was retracted and an auditory tone (4 beeps of 4 kHz; 50-ms duration with 25-ms intervals) signalled the start of the 2-s response period. Based on this structure we divided the trial time in four windows: cue (1.1-2.3 s), touch (2.4-3.3 s), late touch (3.4-4.9 s) and outcome (5-7 s). A water reward was given when the mouse licked in the outcome window after the presentation of the go texture (‘Hit’). The first lick during the outcome window triggered the feedback. Licks during the late-touch window were ignored. A white noise punishment was given for licking in the outcome window for the no-go texture (‘False alarm’, FA). When the mouse withheld licking after the presentation of the go texture (‘Miss’) or the no-go texture (‘Correct rejection’) neither reward nor punishment were given. In the first training session, the identities of go and no-go textures were randomly assigned to the animal and maintained for the whole experiment (go texture: P100 in 3 mice and P1200 in 5 mice).

Animals were kept on a reversed light/dark cycle. After accustoming the mice to the experimenter, habituation to head-immobilization began. We increased head-restraining time with every training session, carrying out two session per day. Mice were water scheduled for behavioural training once they sat quietly for >2 min and were introduced to the experimental setup. Weight, health and water intake were monitored daily. During the first two sessions in the setup, mice only received water reward (∼5 µl per repetition). In session 3 and 4 the go-texture presentation was introduced and an automatic water reward was given, to form an association between texture and reward. Once the mice were able to trigger the water reward autonomously, the no-go texture was introduced starting from presentation in 1% of the cases and gradually increasing to 50%. The first imaging session was scheduled when mice licked consistently for both the textures. Imaging sessions were carried out once per day per animal and lasted as long as a mouse actively engaged in the task (63-209 trials per session). For the first 3-5 imaging sessions go and no-go textures were presented each in 50% of the trials. Thereafter, to facilitate learning, presentation of the no-go texture was repeated in trials following an error trial (false alarm or miss). This ‘repeat-incorrect’ strategy was accounted for in the calculation of behavioral performance by considering the occurrence of the go-texture in a sliding window of 5 trials. Mice learned to differentiate the textures and showed stable expert performance (>75% correct trails) after 12-18 sessions. The performance of each animal (correct response probability as a function of trial ID) was quantified by a state-space smoothing algorithm that provides a learning curve with confidence intervals^65^. The first expert trial and the last naïve trial were identified by an expectation maximization algorithm using a Gaussian state equation. Learning onset (i.e., the last naïve trial) was defined as the trial when the lower 95% confidence interval exceeded 50% correct responses. The first expert trial was defined as the trial, from which on onwards the performance of the animal exceeded chance level with 95% confidence. For analysis of the learning process of all mice, we aligned learning curves to the first expert trial and used a time window of 500 trials before and 150 trials after the first expert trial (trial ID −500 to 150).

### Recording of licking and whisking behavior

Using a 950-nm infrared LED, whisker motion was imaged during the trial at 40 Hz using a high-speed CMOS camera (A504k, Basler). The average whisking angle across all whiskers was analyzed from the videos using a whisker tracking software^66^. The whisker envelope was extracted as the difference between the maximum and minimum whisker angle using the MATLAB function *envelope*. The estimated time point of the first touch between whisker and texture was obtained by calculating the time of the average whisker envelope maximum within the pre-touch and touch window across one session. Licking was estimated based on the event rate from the capacitive lick sensor sampled at 100 Hz. The lick rate was calculated based on the number of lick events in a 200-ms sliding window, assuming that an average lick event lasts 4 ms. Licking and whisking onsets in Hit trials were determined using the MATLAB function findechangepts detecting changes in the root mean square level of the signal. Unrewarded licks occurred prior to the outcome window (1s to 4s in trial time), while rewarded licks occurred within the first half the outcome window (4.8s to 6s). Active touch related whisking onsets occurred during the sensory window (2.3s - 3.5s), while free whisking onsets occurred before this period (1s to 2s).

### Two-photon calcium imaging

In vivo awake calcium imaging was performed using a custom-built two-photon microscope equipped with a Ti:sapphire laser system (Chameleon Ultra, Coherent), a water-immersion objective (CFI LWD 16X/, 0.8 NA; Olympus), a custom-built scanner unit with a 4-kHz resonance scan mirror (CRS 4KHz, Cambridge Technology) and a galvometric mirror (6220H, Cambridge Technology), a Pockel’s Cell (Model 350-80-LA-02, Conoptics, Danbury, CT) and a hybrid-photodetector (HPDs, R11322U-40 MOD, Hamamatsu). The microscope was controlled by the custom-written software Scope^31^ (http://sourceforge.net). An electrically tunable lens (ETL; Optotune EL-10-30-TC, Optotune AG, Zurich, CH; with an plano-concave offset lens, f = −100 mm, Qioptiq) was imaged on the scan mirrors using a 1:1 telescope of f= 100 mm lenses (AC254-100-B-ML, Thorlabs). For initial identification of GCaMP6f-positive neurons, a volume stack was acquired using 800-nm excitation and a green emission filter (510 ± 42 nm bandpass). For calcium imaging, GCaMP6f was excited at 920 nm. Four imaging planes were identified per animal, spanning from close to the pia mater to below the nexus of L5 tufts (approx. −30 µm to −370 µm). Plane hopping was implemented with the ETL ^67^ and images were acquired at 10 Hz volume rate with 508×168 pixel resolution resulting in a 230 µm x 230 µm field of view. Laser power was adjusted per plane ranging from 10 to 65 mW under the objective. Single trials of >7-s duration were recorded with 4-s inter-trial intervals.

### Optogenetic silencing

To transiently suppress apical activity of L5 pyramidal neurons in barrel cortex during reward delivery (in the outcome window), we expressed the outward proton pump eArchT3.0^68^ in S1→S2 L5 neurons using the same surgical procedure of virus injection and window implantation as described for the calcium imaging experiments. In five mice, we made three injections of undiluted AAVretro-hSyn1-chI-FLEX-mCherry_2A_NLS_FLPo virus solution (6.3 × 10^12^ vg/ml) into S2 and three injections of AAV-1/2-hSyn-chl-dFRT-eArchT3.0_EYFP-dFRT virus solution (5.3 × 10^12^ vg/ml) in S1 barrel cortex (coordinates and volumes as described above). In three additional mice we induced expression of eYFP (AAV5-EF1a-fDIO-EYFP_WPRE, 4.9 × 10^12^ vg/ml) instead of eArchT3.0 for control. After the implantation of the glass window, a ferrule holding an optical fiber (910 µm) was positioned and secured in place with dental cement above the window centered over barrel cortex. Animal handling and training was carried out as described above. Once mice reliably licked for water in the experimental setup, 561-nm green laser light (5 mW, CW laser Coherent OBIS-561-50 LS) was delivered through the optical fiber in 100% of the trials. The perturbation only occurred during the outcome window of the trial, lasting 2.4 s (4.8 - 7.2 s in trial time). After 1800 trials of laser perturbation, optogenetic silencing was stopped. The optical fiber transmitting the laser light to the behavioral setup was detached from the ferrule placed above the craniotomy. This change preserved similar light conditions and allowed the mouse to behave and learn without the optogenetic manipulation. Experiments were stopped after 8 weeks of experimentation in accordance with our animal licence. For mice that did not reach expert performance levels, but performed above chance levels at this time, the last recorded trial was considered as their first expert trial. Licking behavior was constantly recorded during the whole experiment. To determine the effect of optogenetic perturbation on whisking behavior, we connected the optical fiber to the ferrule in expert mice and applied laser illumination in 50% of trials. We analyzed whisking behavior as described above and compared the conditions with and without illumination.

### *In vivo* electrophysiological recordings

To validate the effect of optogenetic perturbation we performed acute *in vivo* recordings in lightly anesthetized mice (n = 3) expressing eArchT3.0 selectively in S1→S2 L5 neurons. At the start of validation experiments, animals were anesthetised with isoflurane (2% for induction and <1.5% during recording), and their body temperature was maintained at 37°C using a heating pad. A small craniotomy (<1 mm diameter) was performed over the area of virus injection in barrel cortex and the brain was covered with silicon oil. A silver wire was placed in contact with the cerebrospinal fluid through a small (0.5 mm) trepanation over the cerebellum to serve as reference electrode. A silicon probe (Atlas Neurotechnologies, 32-contact linear array with 50 µm inter-contact spacing) was inserted into the left cortical hemisphere. The top-most electrode was left in contact with the surface of the brain under visual guidance, to ensure that the probe covered the entire cortical column including superficial L1. We used two approaches for optogenetic light illumination. In a first set of experiments, a fiber-optic cannula was positioned to deliver laser light (561 nm, 5 mW) to the surface of the brain just adjacent to the silicon probe, but not inserted into the brain, comparable to the awake in vivo experiments. In a second set of experiments, we opened the skull lateral to barrel cortex and inserted two thin optical fibers (400 µm) into the brain parallel to the brain surface, so that the fiber tips were positioned in cortical L1 and L5. After positioning of the silicon probe and cannula, the preparation was left for 30 min to allow the brain and electrode to stabilise. After stabilisation, the broadband voltage was amplified and digitally sampled at a rate of 30 kHz using a commercial extracellular recording system (RHD2000, Intan Technologies). Spontaneous activity was recorded over 1-1.5h long recording sessions divided into trials (7-s duration, laser on for 2 s) separated by 1-s inter-trial intervals, mimicking the awake optogenetic experiments. The raw voltage traces were processed offline using fourth-order Butterworth filters to separate the local field potential (< 400 Hz lowpass filter) and the multi-unit activity (MUA; bandpass filter 0.46-6 kHz). Subsequently, the local field potential was used to compute the current source density to localize currents arising from the optogenetic stimulation. The high-pass data was thresholded at 5.5 times the standard deviation across the recording session and the numbers of spikes in windows of interest were counted. To combine data across mice, the activity at sites with clear MUA was expressed in percent of the baseline value, i.e. the average spike rate during the period without laser illumination.

### Cleared tissue light-sheet microscopy

Two mice were injected with retrograde AAVretro-hSyn1-chI-FLEX-mCherry_2A_NLS_FLPo virus in S2 and AAV5-EF1a-fDIO-EYFP_WPRE virus in S1 (for details see above) and their brains were cleared using the CLARITY protocol^69,70^. In brief, after 4 weeks of expression, mice were perfused and the brains post-fixed for 48 hours in a hydrogel solution (1% paraformaldehyde, 4% acrylamide, 0.05% bis-acrylamide, 0.25% VA044) before the hydrogel polymerization was induced at 37°C. Then the brains were placed in 40 ml of 8% SDS at room temperature (RT) for approx. 25 days. The brains were put into a refractive index matching solution (RIMS) and equilibrated for 1 day before imaging.

We visualized sparsely GCaMP6f-labeled S1→S2 L5 neurons after clearing brain hemispheres (n = 2) with a custom iDISCO protocol^71^. After 4 weeks of expression, mice were perfused and the brains post-fixed in 4% PFA in PBS for 4.5 hours at 4°C, shaking at 40 rpm. Brain hemispheres were washed in PBS for 3 days at RT and 40 rpm, with daily solution exchange. Samples were dehydrated in serial incubations of 20%, 40%, 60%, 80% methanol (MeOH) in ddH_2_O, followed by 2 times 100% MeOH, each for 1 hour at RT and 40 rpm. Pre-clearing was performed in 33% MeOH in dichloromethane (DCM) overnight (o.n.) at RT and 40 rpm. After 2 times washing in 100% MeOH each for 1 hour at RT and then 4°C at 40 rpm, bleaching was performed in 5% hydrogen peroxide in MeOH for 20 hours at 4°C and 40 rpm. Samples were rehydrated in serial incubations of 80%, 60%, 40%, and 20% MeOH in in ddH_2_O, followed by PBS, each for 1 hour at RT and 40 rpm. Permeabilization was performed by incubating the mouse hemispheres 2 times in 0.2% TritonX-100 in PBS, each for 1 hour at RT and 40 rpm, followed by incubation in 0.2% TritonX-100 + 10% dimethyl sulfoxide (DMSO) + 2.3% glycine + 0.1% sodium azide (NaN_3_) in PBS for 3 days at 37°C and 65 rpm. Blocking was performed in 0.2% Tween-20 + 0.1% heparine (10 mg/ml) + 5% DMSO + 6% donkey serum in PBS for 2 days at 37°C and 65 rpm. Samples were stained gradually with primary polyclonal chicken-anti-GFP antibody (Aves Labs, GFP-1020) and secondary donkey-anti-chicken-AlexaFluor488 antibody (Jackson ImmunoResearch, 703-545-155) 1:400 in 0.2% Tween-20 + 0.1% heparine + 5% DMSO + 0.1% NaN_3_ in PBS (staining buffer) in a total volume of 1.5 ml per sample every week for 4 weeks at 37°C and 65 rpm. Washing steps were performed in staining buffer 5 times each for 1 hour, and then for 2 days at RT and 40 rpm. Clearing was started by dehydrating the samples in serial MeOH incubations as described above. Delipidation was performed in 33% MeOH in DCM o.n. at RT and 40 rpm, followed by 2 times 100% DCM each for 30 minutes at RT and 40 rpm. Refractive index (RI) matching was achieved in dibenzyl ether (DBE, RI = 1.56) for 4 hours at RT.

3D stacks of cleared brains and hemispheres were acquired using a mesoSPIM light-sheet microscope^72^ (www.mesospim.org). Imaging data were post-processed using custom-written routines in MATLAB. To visualize neurons, local contrast enhancement was performed per slice by subtracting a Gaussian-smoothed version of the slice (4σ). Barrels were visible in the green autofluorescence channel. An anatomical barrel map was fitted to the barrel autofluorescence using the MATLAB functions *cpselect* and *fitgotrans*. 3D volume projection was performed using Imaris (9.8.0, Oxford Instruments).

### Confocal histology

After the last awake imaging session mice were administered a lethal dose of pentobarbital (Ekonarcon, Streuli) and transcardially perfused with sterile NaCl (0.9%) followed by 4% paraformaldehyde (PFA, 0.1 M phosphate buffer, pH 7.4). From 100-µm thick coronal brain slices we acquired histological images with a confocal laser-scanning microscope (Olympus FV1000). Coronal sections were registered to Paxinos and Franklin’s mouse brain atlas using manually set landmarks using *cpselect* (MATLAB) and aligning the atlas via *fitgeotrans* (MATLAB).

### Morphological reconstructions

Anatomical two-photon image stacks of all fields of view were acquired before behavioral training using the two-photon microscope with 800-nm laser excitation. 3D reconstructions of imaged dendritic tufts were obtained using the semi-manual interpolation option of the VolumeSegmenter app in MATLAB. Tuft membership was determined based on the morphological stacks as well as on high correlation of calcium signals. In 42 neurons, the trunk could be clearly identified together with its corresponding daughter tuft branches. The comparison of functional class within dendritic tufts as well as the LE/GE analysis were performed on this subset of neurons.

### Preprocessing and visualization of calcium imaging data

Motion correction of the acquired movies of GCaMP6f fluorescence was carried out by a custom-written Python pipeline using the NoRMCorre algorithm for non-rigid artefact correction provided by CaImAn^73^. Single non-overlapping dendritic branches were identified and regions of interest (ROIs) were defined manually for each session. A consistent nomenclature was used to identify the same dendritic branches over consecutive sessions. If multiple ROIs along the same branch were identified, only the ROI closest to the trunk was used for further analysis. Calcium indicator fluorescence signals were extracted using custom software routines written in MATLAB (Mathworks). Background fluorescence was estimated in a background ROI as the bottom 1^st^ percentile fluorescence signal across the entire session and subtracted before calculating the relative percentage change of fluorescence from baseline ΔF/F = (F-F_0_)/F_0_. Baseline fluorescence F_0_ was computed as 51^st^ percentile of the fluorescence signal in a 4-s sliding window. ΔF/F traces were smoothed with a 5-point 1^st^-order Savitsky-Golay filter. Upon visual inspection, we manually excluded calcium traces with obvious artefacts such as motion-induced artefacts, light reflections from the texture, or non-physiological calcium traces. All remaining data were visualized using an Euclidian-distance based UMAP embedding (UMAP embedding 1; Supplementary Fig. 6) with a neighborhood size of 100 data points in the high-dimensional space. For further analyses, detectable transients were defined as fluorescence signals that deviated from baseline noise by >5.5 standard deviations. Noise levels per ROI were determined as the median of the absolute value of the first derivative of the concatenated ΔF/F trace of one session. The set of trials with detectable ΔF/F transients was visualized using a correlation metric-based UMAP embedding (UMAP embedding 2) and a neighborhood size of 30 data points in the high-dimensional space. Analysis and data exploration was carried out using dataspace^30^ (https://github.com/skollmor/dspace) and custom-written MATLAB code.

To determine the onsets of calcium transients, we identified the highest peak in a given trial (*findpeaks* MATLAB function; 1-s minimal distance between peaks and >25% ΔF/F peak prominence). Local maxima were not included in further analysis. For the calcium transient with maximal peak, the onset time point was defined as the minimum of the first derivative of the ΔF/F trace up to 1 s prior to the detected peak. Transients with their peak position within the start window (0-1 s of trial time) were not considered as their onset likely occurred during the inter-trial interval.

### Functional co-evolution of dendritic signals and transformation clustering

To assess learning-related functional changes of trial-related calcium traces in individual dendritic branches or trunk ROIs we employed the custom-developed approach of “transformation clustering” that is inspired by earlier work using nearest neighbour graphs to understand high dimensional data^30^. The dimensionality of the raw ΔF/F trace vectors (70 frames) was reduced to 12 dimensions using multi-dimensional scaling (MDS). Transformation clustering exclusively used this 12-dimensional representation of ΔF/F traces. It is independent from the dimensionality-reduced UMAP embedding, which is employed to visualize the data set. Functional changes across the learning time course in one dendritic branch were compared to the changes of any other dendritic branch in our dataset as follows: Let *d* and *d*’ denote a pair of dendritic ROIs with trial responses 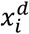 and 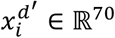 (7-s recording at 10 Hz), where *i* denotes the index of the trial (the trial ID) in which the response was recorded. Note that many trials contain no detectable transients (as defined above). Let *i*_1_, …, *i*_*k*_ denote the indices of the trials in which dendritic ROI *d* shows a detectable transient. Let *NN*^*k*^(*d*, *i*, *d*′) denote the set of *k* nearest neighbours of the trial response 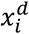 among all responses 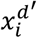 for dendritic ROI *d*′ that show detectable transients. Let 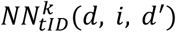 denote the set of trial indices of the nearest neighbours of the trial response 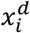 among the activations for ROI *d*^′^:

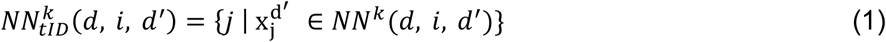

Finally, let 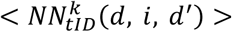 denote the average of those trial indices. We define the similarity of *d* and *d*′ as the correlation coefficient between the two vectors [*i*_1_, …, *i*_*k*_] and 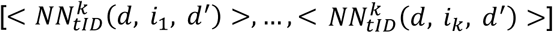 We refer to this correlation coefficient as *Tcc*_*dd*_′ (transformation correlation coefficient). To assess significance of the correlation, we compared the actual *Tcc*_*dd*_′ to a null hypothesis derived by shuffling trial IDs for dendritic ROI *d’*. Shuffling removes the long-term temporal relationship between ROIs *d* and *d*′. We define the corrected transformation correlation coefficient, *CT*_*dd*_′, as the inverse percentile of *Tcc*_*dd*_′ with respect to the null hypothesis distribution generated by many shuffle iterations (e.g. 1000).

Let **CT** denote the square matrix of corrected transformation correlation coefficients for all pairs of dendritic ROIs *d* and *d’.* We compute a symmetric dissimilarity matrix, **D**, through **D** = −(**CT** + **CT**^T^), where **CT**^T^ denotes the matrix transpose of **CT**. We applied hierarchical link (HL) clustering to extract transformation clusters. We achieved comparable results using k-means clustering after applying multidimensional scaling to the dissimilarity matrix **D**. Additionally, we employed fuzzy c-means clustering to obtain a gradual measure of how likely a dendrite belongs to one of the two binary clusters assigned by HL. For fuzzy c-clustering we used an exponent for the partition matrix of 1.3 to reduce overlap between the two clusters and qualitatively replicate the HL clustering results.

Note that only dendritic ROIs with more than 40 ΔF/F traces with detectable calcium transients spanning a range of more than 500 trial IDs were included in the initial transformation clustering analysis. The results from this initial clustering were greedily expanded in order to assign cluster IDs also to dendritic branches and apical trunks that had only 6-39 trials with detectable calcium transients (independent of the range of recorded trial IDs). For these dendritic ROIs we calculated corrected transformation correlation coefficients with all other previously classified dendritic ROIs. The 5 previously classified ROIs with highest coefficients values (corresponding to the smallest dissimilarities) were selected and the predominant cluster ID of this set was as assigned to the unclassified ROI.

### Dendritic event detection

For every tuft and trial, we summed ΔF/F for all tuft ROIs and identified candidate events as local maxima separated by at least 2 s (*findpeaks* function, MATLAB). For each candidate event we generated a ΔF/F heatmap for a 2-s window around the detected peak (matrix with ROIs as rows and 20 ΔF/F time points as columns), representing the activity of trunk and tuft branches in individual L5 neurons. For identification of local events we defined two metrics. First, we defined trunk-tuft ‘co-activity’ as the correlation of the ΔF/F traces in the trunk and the tuft branch *tuft*_max_ with the highest ΔF/F amplitude (defined as min-to-max range). Second, we defined ‘apicality’ as 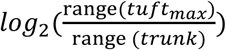. With these definitions, local events had to fulfil three criteria: (1) the maximum z-scored ΔF/F for at least one ROI exceeds 2 (z-scoring is performed with respect to all trials recorded for this ROI); (2) co-activity is negative or non-significant (p > 0.05, Pearson’s correlation, MATLAB); and (3) apicality is larger than 2, i.e., the signal range of the maximal tuft branch activity is at least four times higher than in the trunk.

### Decoder discrimination analysis

We trained three types of linear support vector machine (SVM) decoders from different trial types: Hit vs. CR (different textures and actions), Hit vs. FA (different textures, same actions), and CR vs. FA (same textures, different actions). For each trial type combination, decoders were trained using the pooled activity from either sensory or outcome window, using either whole-tuft activity or single dendrite activity. A separate decoder was trained for each imaging session. For each session, we randomly divided all the matched pairing trials into 5 subsets. We trained 5 decoders by excluding one subset at one time, therefore each decoder was trained on 90% of the training set. To avoid overfitting, we regularized the SVM coefficients with ridge (L2) penalty. The regularization term was cross-validated in a log space of 10 parameters from 10-5 to 101. To avoid overfitting due to unbalanced class number, we also implemented a misclassification cost that is inversely correlated with the total number of each class. To evaluate the decoder performance, the ROC (receiver operating characteristic) curve and the AUC (area under curve) were calculated. For null distribution, we randomly permuted the neuron indices in the activity matrix, and applied the trained decoders to obtain shuffled AUC values. We then tested the AUC from real data against 5% or 95% quantiles of the shuffled distributions.

### TD error-based computational model

We replicated our experimental results in a temporal difference (TD) reinforcement learning model. In brief, an estimator *V̂* of the state-value function *V* was learned using the TD error^5^

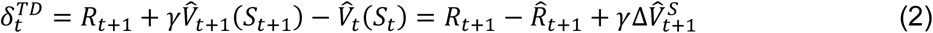

where *R*_*t*+1_is the reward outcome at time *t* + 1, *S*_*t*_ the state at time *t*, −*R̂*_*t*_ represents the reset in *V̂* when an outcome (good or bad) is expected, 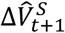 represents the change in state-value estimation associated with the state change from *S*_*t*_ to *S*_*t*+1_, and *γ ∈* [0,1] is a discounting parameter for future rewards (in our model we presumed *γ* = 1, i.e., no discounting; for completeness we include it our desription).). The derivation for this formulation of the TD error is described in the Supplementary Note 1. Our model postulates that apical dendrites convey salience information to the neurons to modulate their somatic output activity and steer behavior according to the animal’s sensory experience and needs. First, we consider ***unconditioned salience*** *s*^*U*^ as the absolute value of unexpected outcome, i.e.,

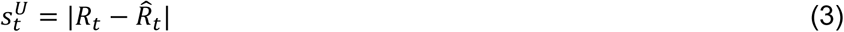

because only the difference between the actual reward *R*_*t*_ at time *t* and the estimated (expected) reward *R̂*_*t*_ is informative for the animal (variables with a hat indicate model estimations). Initially, when the outcome is entirely unexpected, *s*^*U*^ is large; once the task has been learned and the outcome can be predicted, this term approaches zero.

Second, we assign salience to any sensory stimulus that is evaluated to be task-relevant. If a stimulus bears predictive power for future outcome, the perception of this stimulus will also impact the state-value estimation *V̂*. We define the ***conditioned salience*** *s*^*C*^ that develops during learning as the absolute value of the change Δ*V̂*_*t*_ in state-value function estimation when moving from *t* − 1 to *t*:

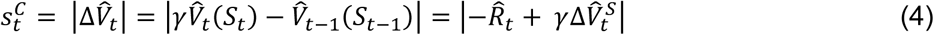

Both variables *s*^*U*^ and *s*^*C*^ are unsigned terms that relate to components of the classical TD error (see Supplementary Note 1 for a detailed derivation).

We implemented these ideas in a network model of rate-based multi-compartment neurons, imposing the two types of salience signals on apical dendritic compartments (Fig. 3b). We designed the simulation to represent a population of L5 pyramidal neurons, with each neuron modeled as three compartments: the soma and two apical dendritic branches: one outcome branch receiving unconditioned salience and one sensory branch receiving conditioned salience information. The somata received bottom-up sensory stimulation via basal inputs, and their activities were gain-modulated by the apical dendrite activity. Sensory stimuli were represented by binary variables that included the mutually exclusive go and no-go texture stimuli (occuring with 50% probability each and with fixed timing in the texture window), the auditory cue stimulus occurring in each trial in the cue window, and additional distractor stimuli, for which variable probabilities of activation were preset and, if they became active, the activation time (once during each trial) was uniformly drawn for each trial. Each simulated trial consisted of 18 time bins (4 Hz sampling) starting with one time bin before the tone-cue, covering cue period (1 s), touch period (1 s), and late-touch period (2 s), and ending with one outcome time bin. In the pure-selectivity model, the bottom-up basal inputs **X**^bas^ were connected one-to-one to the L5 somata, producing one go-texture-responsive (Go) neuron, one no-go-texture-responsive (NoGo) neuron, and one tone-cue-responsive (Cue) neuron. In addition, we generated 200 distractor neurons whose activities were randomly assigned according to Bernoulli distributions (Supplementary Fig. 12a). We also explored a mixed-selectivity model, in which the input stimuli for go-texture, no-go-texture, tone-cue and 14 distractors were probabilistically connected to the somata of the L5 neurons (Supplementary Fig. 14a). In both the pure-selectivty and the mixed-selectivity model, the bottom-up sensory activation pattern remained fixed throughout the entire simulation (see Supplementary Note 2 for further details).

For each neuron *i*, the activity of the apical dendritic compartment was modeled as the sum of outcome and sensory branch activity

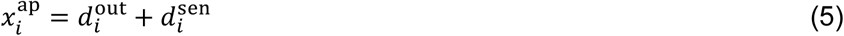

This dendritic activity affected the somatic activities via multiplicative gain modulation:

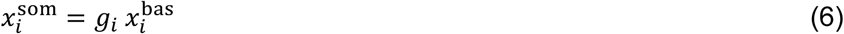

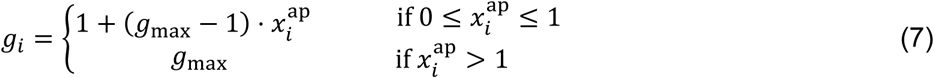

where we chose *g*_max_ = 10. Whereas the unconditioned salience input to the outcome branches was uniform across all neurons, individual neurons learned to adapt the sensitivity of their sensory branch to the conditioned salience input by adjusting the input weights 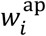. Effectively, this framework learns to amplify the apical gain of task-relevant neurons with *s*^*C*^ = |Δ*V̂*_*t*_|, i.e., in time windows during which relevant information for outcome prediction is available and the state-value estimator *V̂* thus is updated.

To simulate the optogenetic inhibition experiment, we perturbed the model neurons for 1’800 trials by setting the apical dendritic activity **X**^ap^ to zero during the outcome window. For further details of the computational theory and the model implementation see Supplementary Notes.

The code reproducing the simulations and visualisations for the computational model can be found at https://github.com/HelmchenLabSoftware/td_dendrites.

## Supplementary Figures

**Supplementary Fig. 1.**
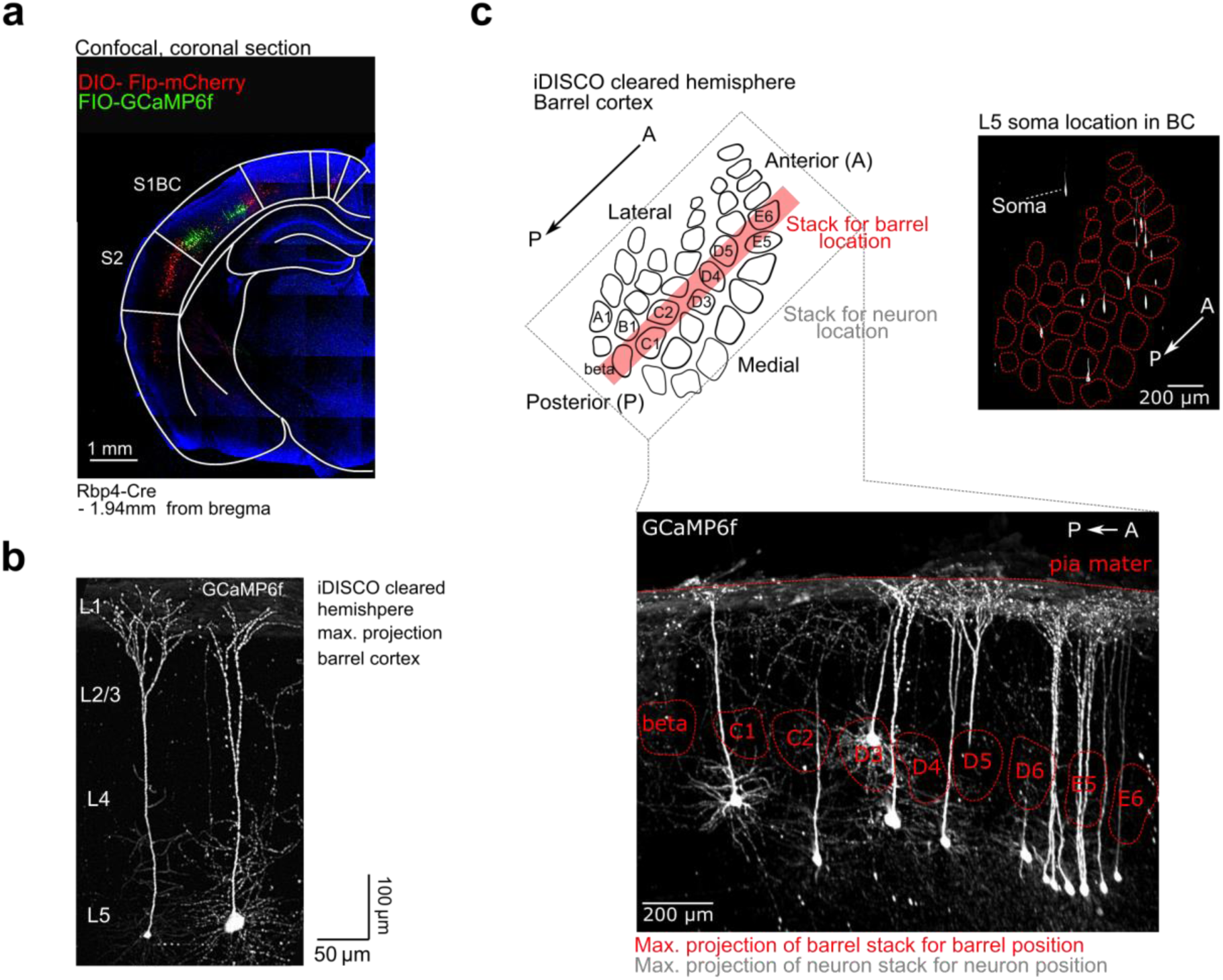
Histological characterization of imaged S1→S2 L5 pyramidal neurons. **a**, Coronal confocal overview image of GCAMP6f-labeled L5 pyramidal neurons in barrel cortex (green) and retrogradely-labeled mCherry-tagged neurons (red) with projections to S2. **b,** Maximum-intensity projection of a confocal image stack of sparsely labeled L5 neurons expressing GCaMP6f in S1 barrel cortex (150 μm slice). Note that most apical dendrites were cut in this slice preparation. **c,** Visualization of the location of sparsely-labeled GCaMP6f-expressing L5 neurons in barrel cortex from an iDISCO-cleared hemisphere. *Top*: Dorsal view of L5 soma location in barrel cortex, with barrel map overlaid according to L4 autofluorescence. *Bottom*: Sagittal view of L5 neurons in barrel cortex along the A/P axis. Note, the maximum-intensity projection of GCaMP6f-expressing neurons was calculated across the whole of barrel cortex (1.3 mm slice) whereas the barrel map outline was derived from a reduced stack (210 μm slice) as visualized in the schematic above (red strip).

**Supplementary Fig. 2.**
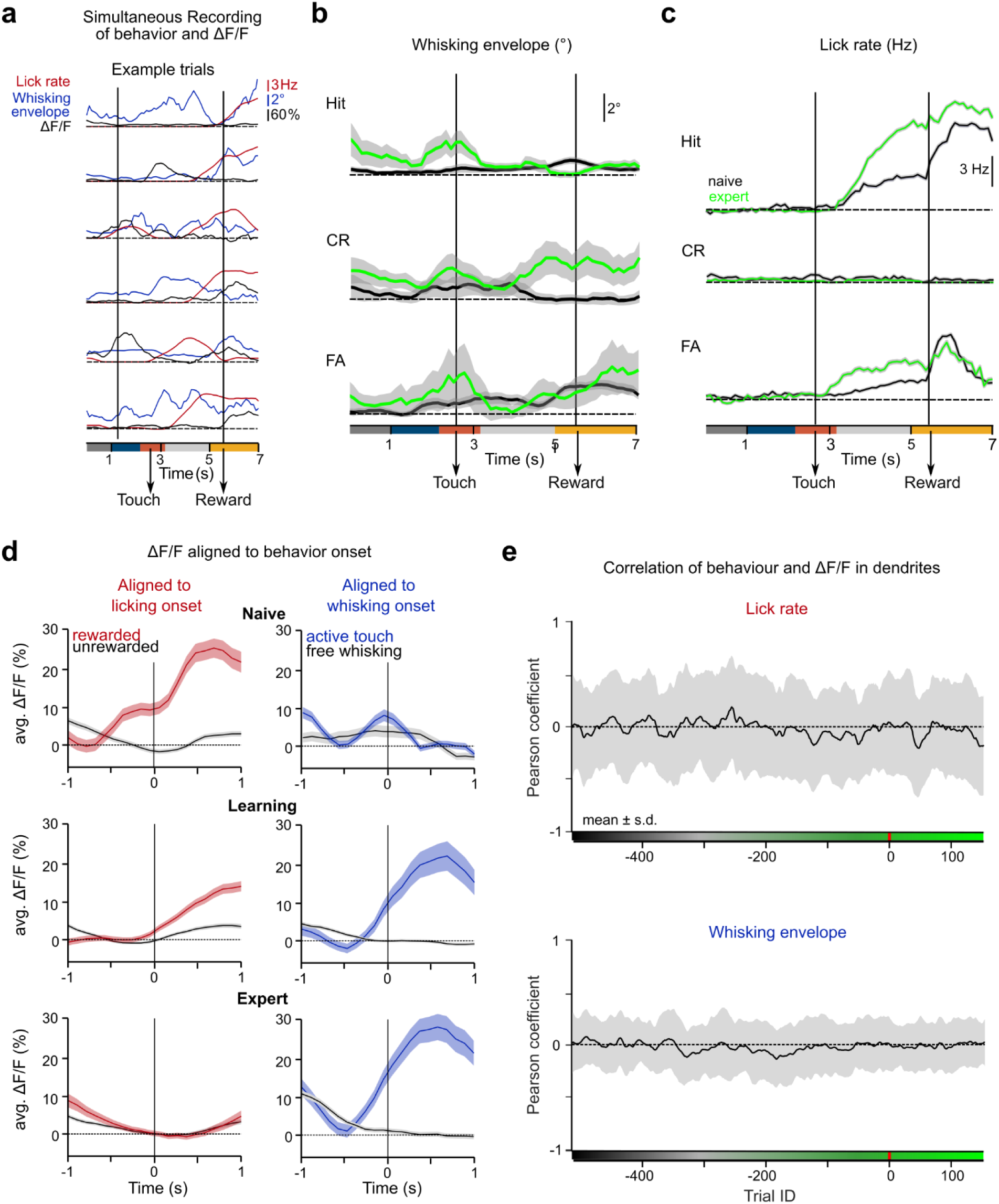
Changes of behavioral variables during learning. **A,** Example traces of lick rate, whisking envelope, and ΔF/F traces of six example trials aligned to trial start. **b,** Average whisking envelope in naïve and expert mice split by trial type (524 naïve Hit, 380 expert Hit, 181 naïve CR, 295 expert CR, 861 naïve FA and 191 expert FA trials, 8 mice, mean ± s.e.m.). The last naïve and the first expert trial per mouse were determined using a state-space model. **c,** Average lick rate in naïve and expert mice split by trial type (same number for trials as in b); 8 mice, mean ± s.e.m.). **d,** Average ΔF/F traces around licking onsets (±1 s) for rewarded and unrewarded licks (left) and around whisking onsets for free whisking and active touch whisking in naïve, learning, and expert condition (right) (Hit trials, mean ± s.e.m.) **e**, Mean correlation of lick rate and dendritic ΔF/F traces (top) and of whisking envelope amplitude and dendritic ΔF/F traces (bottom) across learning. Black line and grey-shaded area indicate mean ± s.d. across all ROIs.

**Supplementary Fig. 3.**
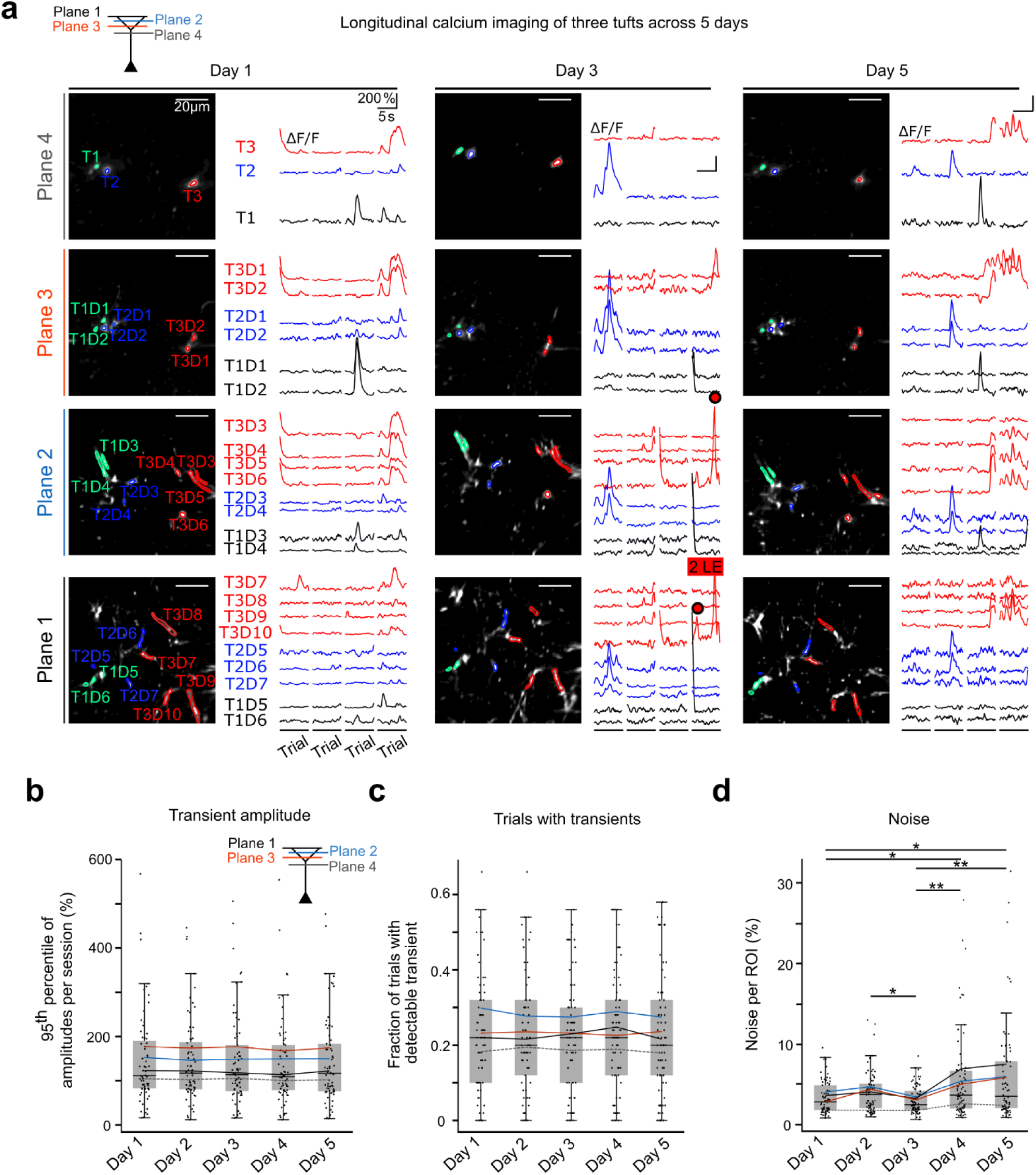
Longitudinal calcium imaging of dendritic activity. **A,** Example two-photon calcium imaging with four imaging planes for three imaging sessions spread over 5 days. The trunks (T) and tuft dendrites (D) of several neurons are shown in different colors. ROIs and respective calcium transients are display for four example trials per session. Two local events (LE) on day 3 are highlighted. **b,** Distribution of calcium transient amplitudes per dendrite (calculated as the 95^th^ percentile of ΔF/F traces with detectable calcium transients in 50 randomly selected trials per day) across five consecutive imaging days. Colored lines indicate average transient amplitude per imaging plane. **c,** Distribution of the number of trials with detectable calcium transients per dendrite. **d,** Distribution of noise levels per ROI calculated based on 50 randomly selected trials per day.

**Supplementary Fig. 4.**
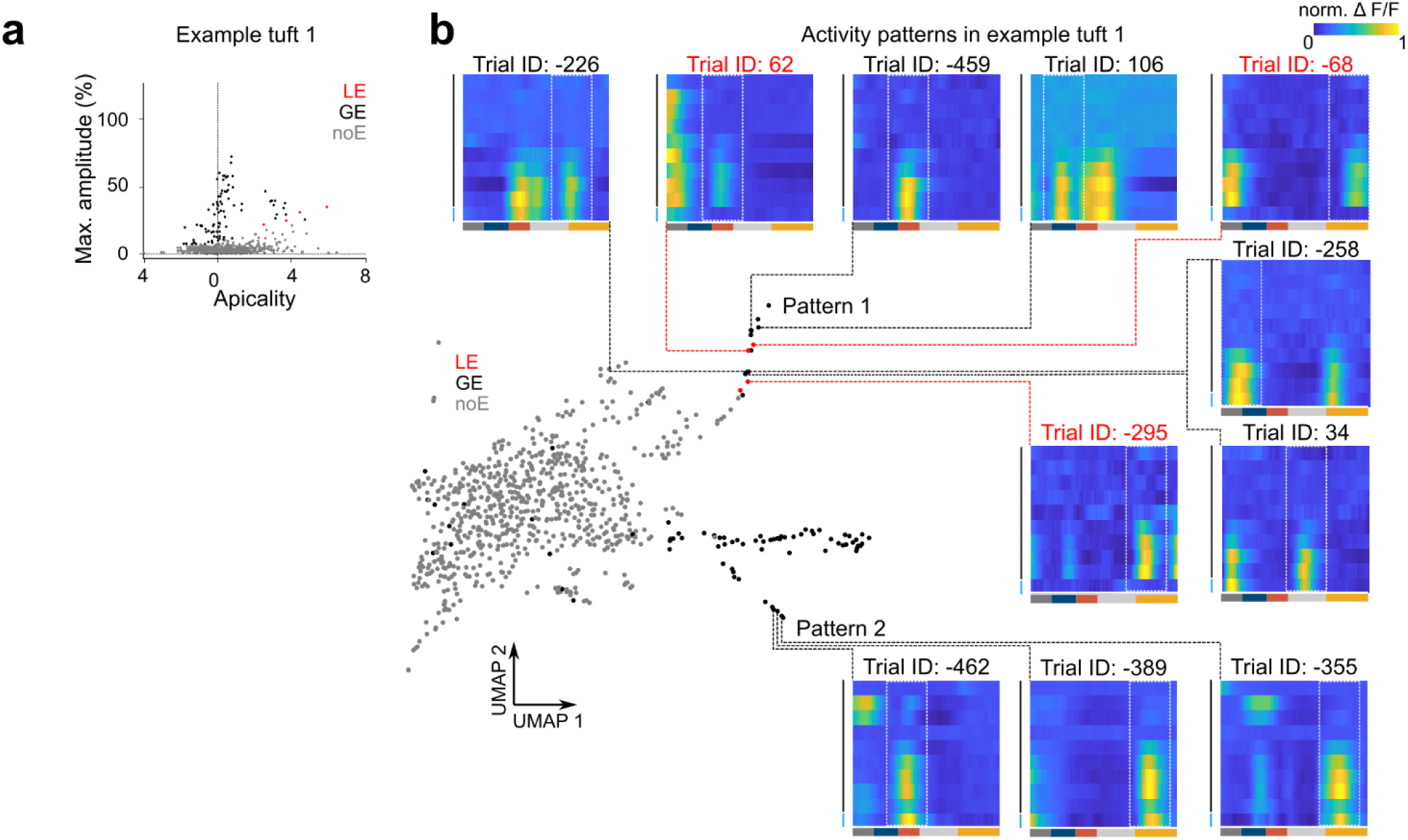
Apical tufts display diverse ΔF/F activity patterns across learning. **A,** Overview of events detected in example tuft 1 colored by event type. **b**, UMAP embedding of all events recorded in example tuft 1 colored by trial ID. Surrounding heatmaps display averaged normalized ΔF/F tuft activity patterns across trail time from selected neighborhoods. Event-centered windows for data analysis are shown by white lines.

**Supplementary Fig. 5.**
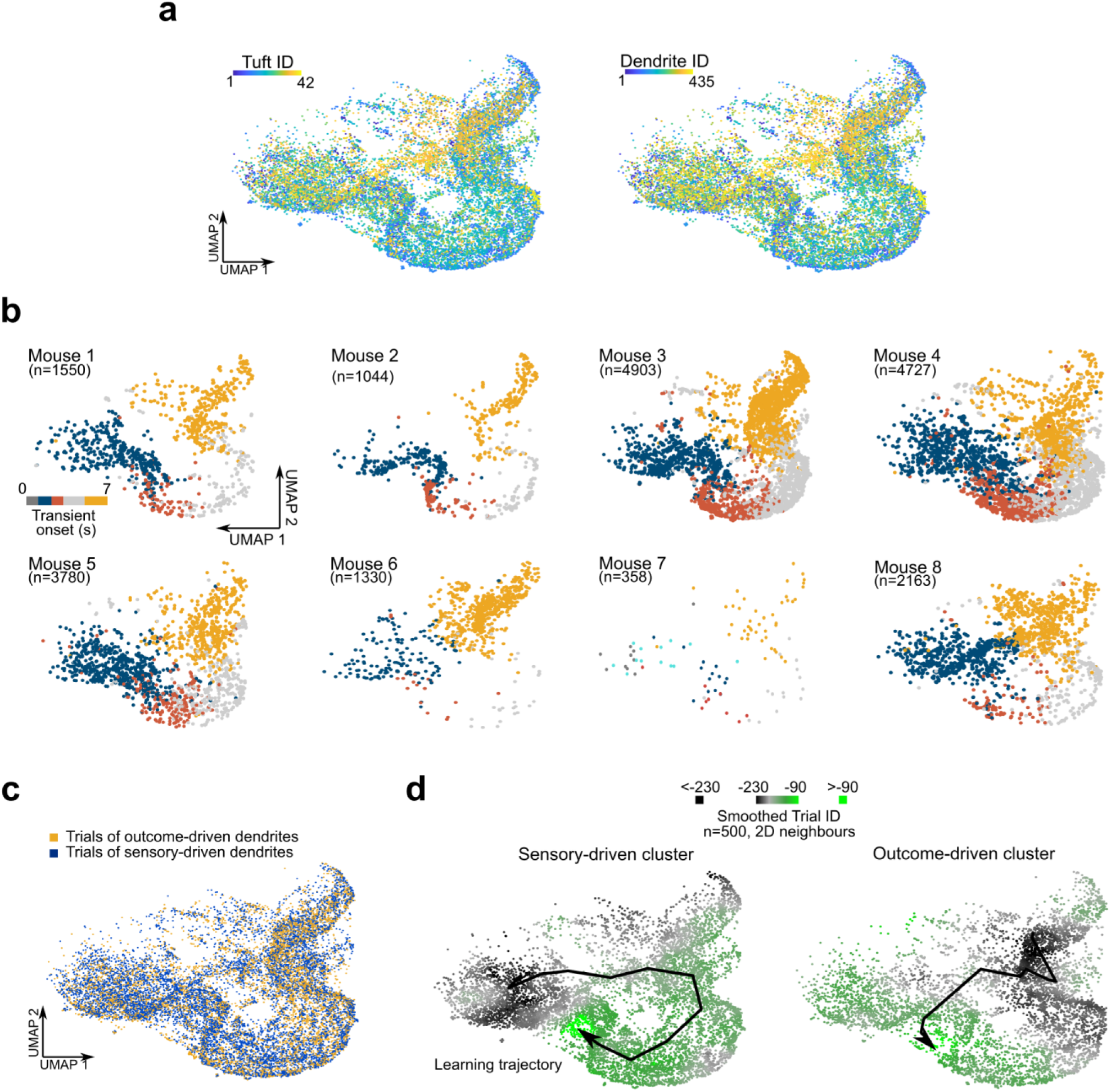
Mapping of various features of trial-related dendritic calcium signals in the UMAP space. **A,** UMAP embedding of all ΔF/F traces with detectable calcium transient onsets within trials colored by mouse ID (left), tuft ID (middle), and dendrite ID (right; 22’027 data points, NN=30). **b**, UMAP embedding separated by mice and color-coded by transient onset time. Value in brackets indicate the number of data points (dendritic ΔF/F traces with detectable transient onsets from single trials) obtained from each mouse. **c**, UMAP embedding of ΔF/F traces of sensory and reward class colored according to their cluster ID. Note that trials of both classes are inter-mixed within the UMAP embedding. **d**, Clusters distinguish themselves by their temporal profile of learning-related changes. UMAP embedding of ΔF/F traces in dendrites of sensory (left) and outcome (right) cluster, colored according to smoothed trial ID. The arrows indicate learning-related trajectories of the centre of mass in the 2D UMAP plot (25-trials binning).

**Supplementary Fig. 6.**
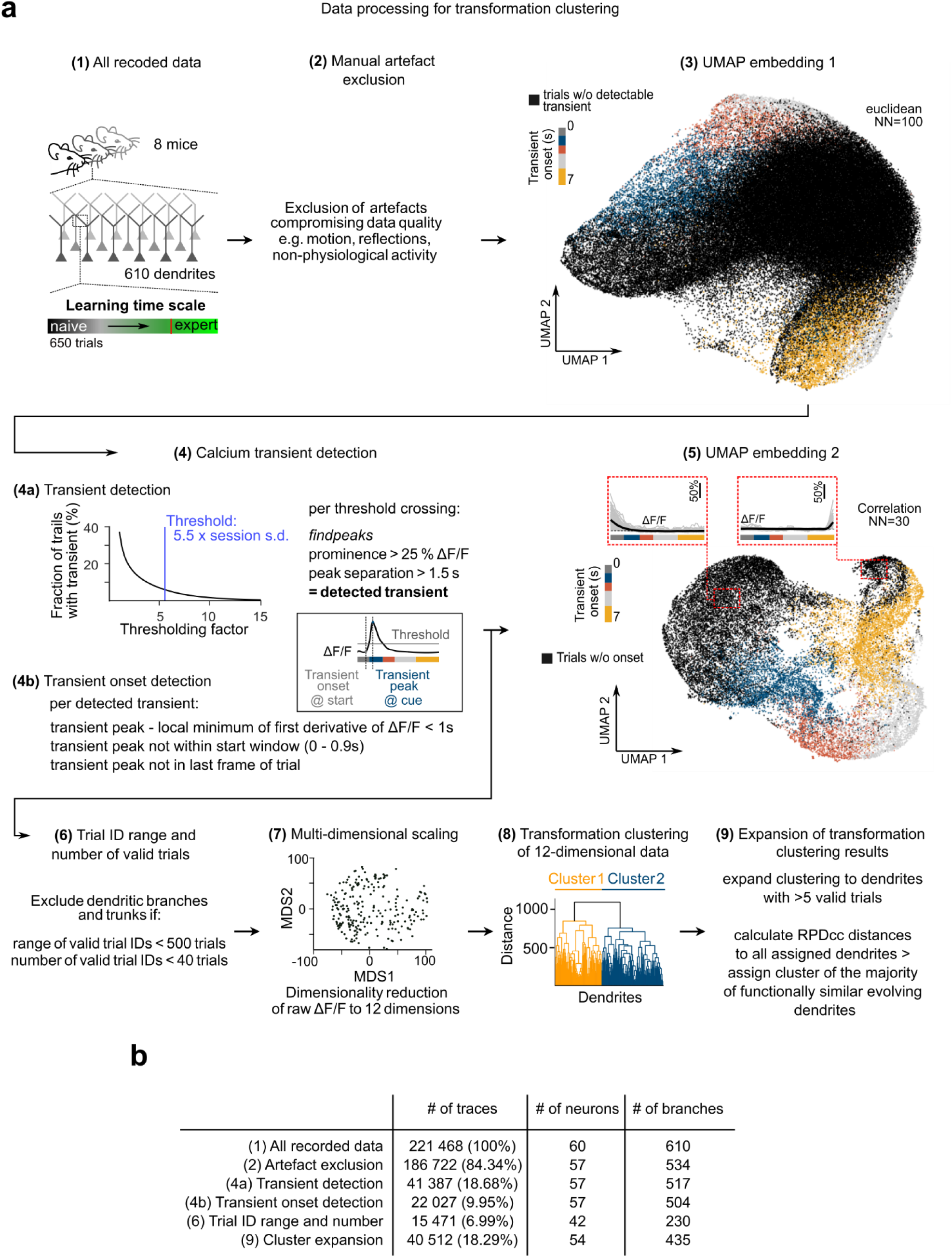
Processing pipeline for dendritic ΔF/F data. **A,** Flow chart for ΔF/F data processing. **b,** Number of traces, neurons and dendritic branches per processing step. Note not for every neuron a trunk was recorded. Color in (3) and (5) are based on transient onset times (4b).

**Supplementary Fig. 7.**
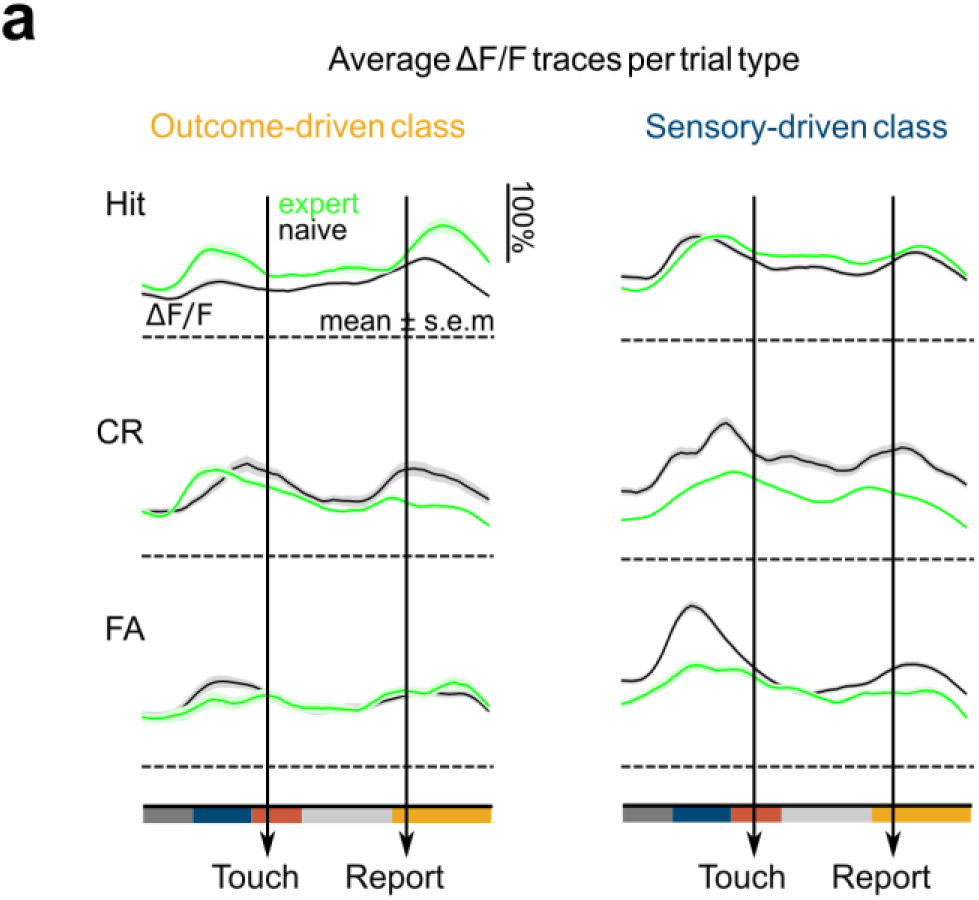
Average ΔF/F traces per trial type and functional cluster. **A,** Average ΔF/F traces across trial types for dendritic branches belonging to reward (left) and sensory (right) class. Averages include only trials of tuft branches and apical trunks with a detectable calcium transient onset and are shown separately for the naïve (black) and expert (green) condition. Note that average ΔF/F traces reflect both transient abundance and transient amplitudes, while transient onset probabilities shown in Fig. 2c,d do not consider amplitudes. (Outcome-driven dendrites: 1452 naïve Hit, 964 expert Hit, 304 naïve CR, 597 expert CR, 1472 naïve FA, 402 expert FA trails; Sensory-driven dendrites: 1383 naïve Hit, 1671 expert Hit, 566 naïve CR, 1283 expert CR,2377 naïve FA, 714 expert FA trails; n = 8 mice; mean ± s.e.m).

**Supplementary Fig. 8.**
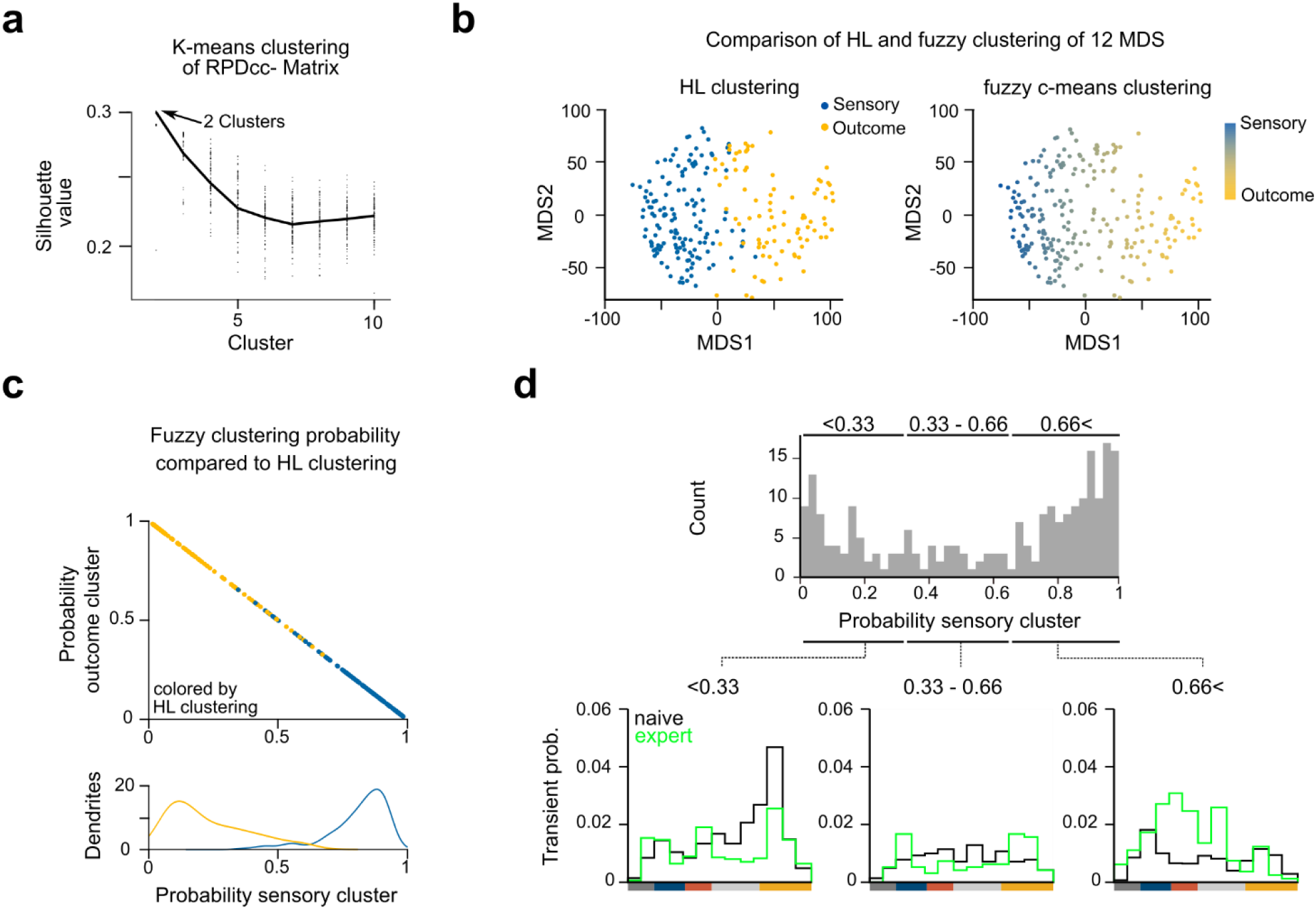
Fuzzy c-means clustering describes a continuum of dendritic cluster identities. **A,** Relationship between silhouette values and cluster numbers obtained as an alternative clustering approach for transformation clustering using k-means clustering. Black line indicates mean silhouette value (k = 2-10 clusters, 100 repetitions each). b, Comparison of hierarchical link clustering results and fuzzy c-means clustering results for two clusters on the first 12 dimensions of multi-dimensional scaling per dendrite. **c,** Top: Relationship of fuzzy c-means clustering probabilities for the sensory- and the outcome-driven cluster, colored according to hierarchal link cluster assignment. Bottom: Distribution of sensory cluster probability across sensory and outcome branch polulation. **d**, Top: Distribution of fuzzy c-mean clustering probabilities for the sensory-driven cluster across all dendrites. Bottom: Distribution of calcium transient onset probability across trial time in the first, middle and last third of the fuzzy c-means cluster probability distribution.

**Supplementary Fig. 9.**
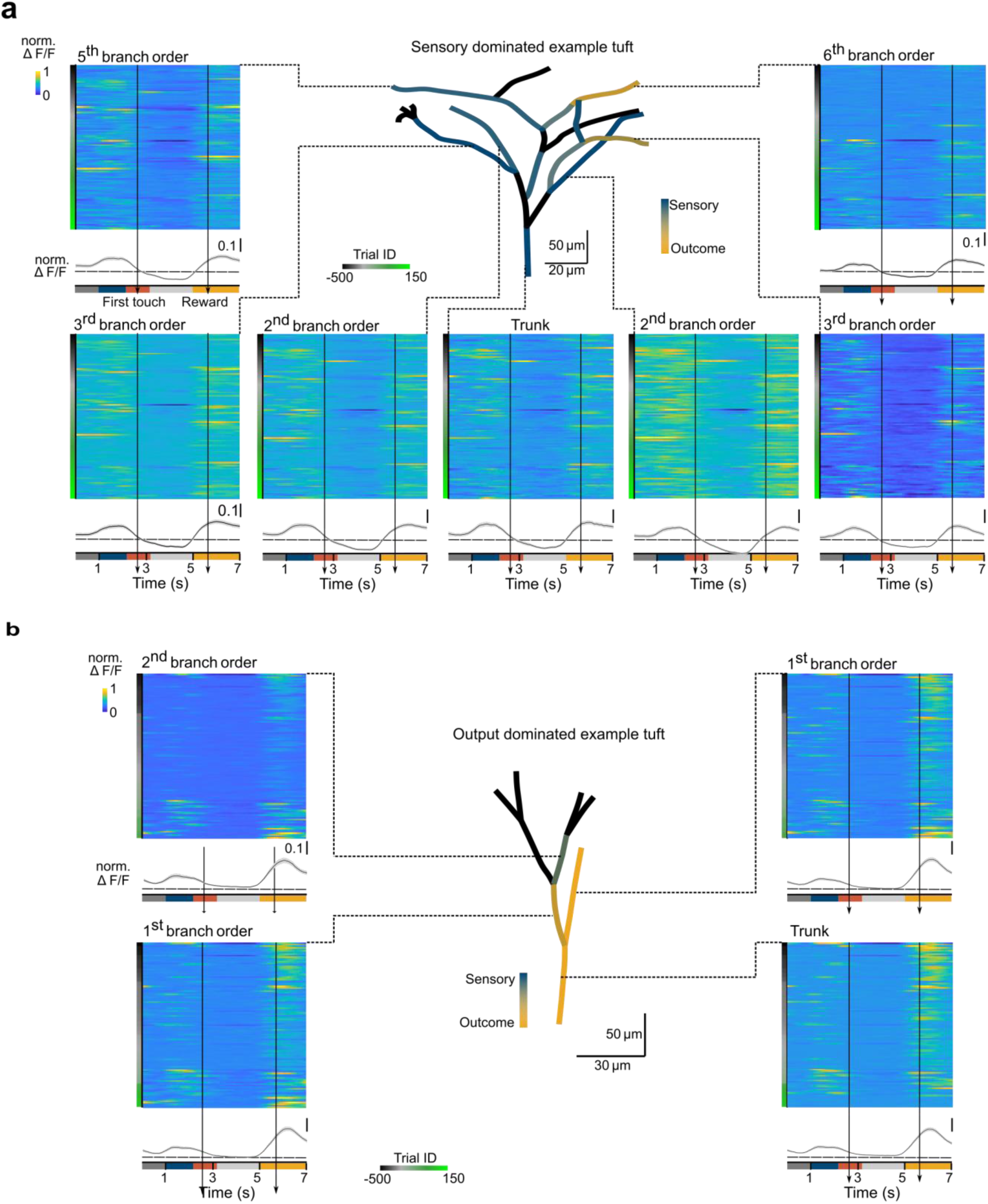
ΔF/F activity changes are distinct in single branches belonging to the same tuft. **A,** Reconstruction of a sensory cluster-dominated example tuft. Single branches are colored based on fuzzy clustering results. Respective heatmaps display normalized ΔF/F activity per branch across learning. Heatmaps include all trials with a detectable calcium transient in the trunk ROI (107 trials). Averaged ΔF/F traces across trial time are displayed underneath the heatmaps (mean ± s.e.m.). **b**, Reconstruction, heatmaps and averaged ΔF/F traces of an outcome cluster-dominated example tuft (112 trials).

**Supplementary Fig. 10.**
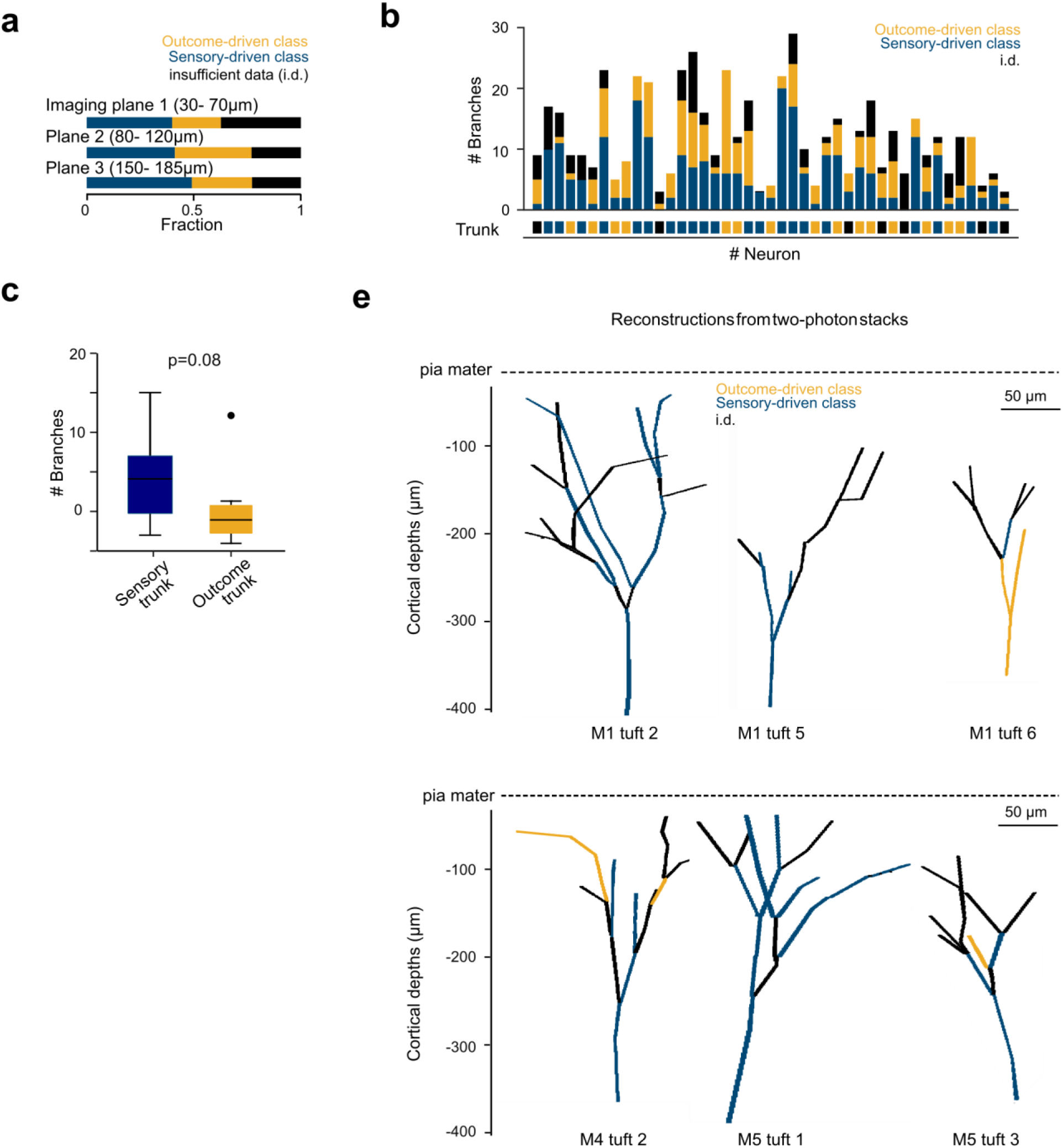
Morphological reconstructions of imaged dendritic tufts and their functional composition. **a**, Fraction of apical branches per class across imaging planes. **b**, Number of branches per functional class per tuft (n = 42). Dendrites that were not assigned to a class due to insufficient data point are shown in black. **c,** Absolute number of apical in sensory and reward tufts (13 ± 7 and 9 ± 5 branches, respectively; mean ± s.d.; p = 0.08, Student t-test; Black line: median, box: 25^th^ and 75^th^ percentile, whiskers: 5^th^ and 95^th^ percentile, outliers: points. **d,** Examples of L5 apical dendritic tufts reconstructed from two-photon image stacks. Branches are color-coded according to their functional class.

**Supplementary Fig. 11.**
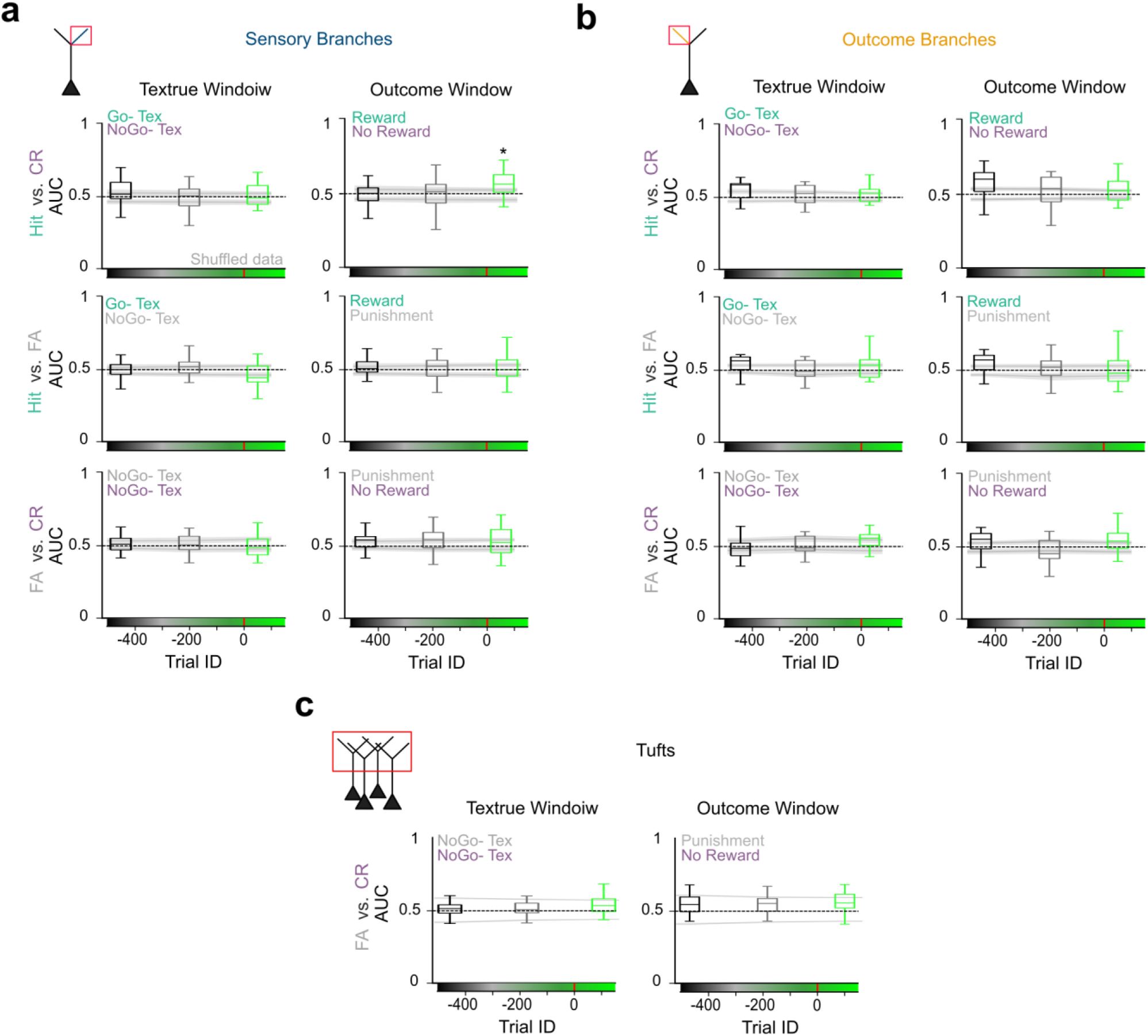
AUC-based discrimination ability of L5 tuft dendritic branches. **a**, Pooled linear SVM discrimination power for ΔF/F activity in Hit vs. CR, Hit vs. FA and CR vs. FA trials of sensory-driven branches across learning. SVM-decoders were trained on the texture window (2.4 - 3.9s) and the outcome window (5 - 7s). Grey lines indicate 5% and 95% quantile of randomly shuffled data. (n=48, 67 and 27 branches for naïve, learning and expert condition; Outcome window, Hit vs. CR: p*=0.04). **b**, Pooled discrimination for outcome-driven branches (n=22, 21, and 25 branches) **c**, Pooled discrimination in all dendritic tufts between FA versus CR trials (n=42 tufts), complementing the plots in Fig. 3a.

**Supplementary Fig. 12.**
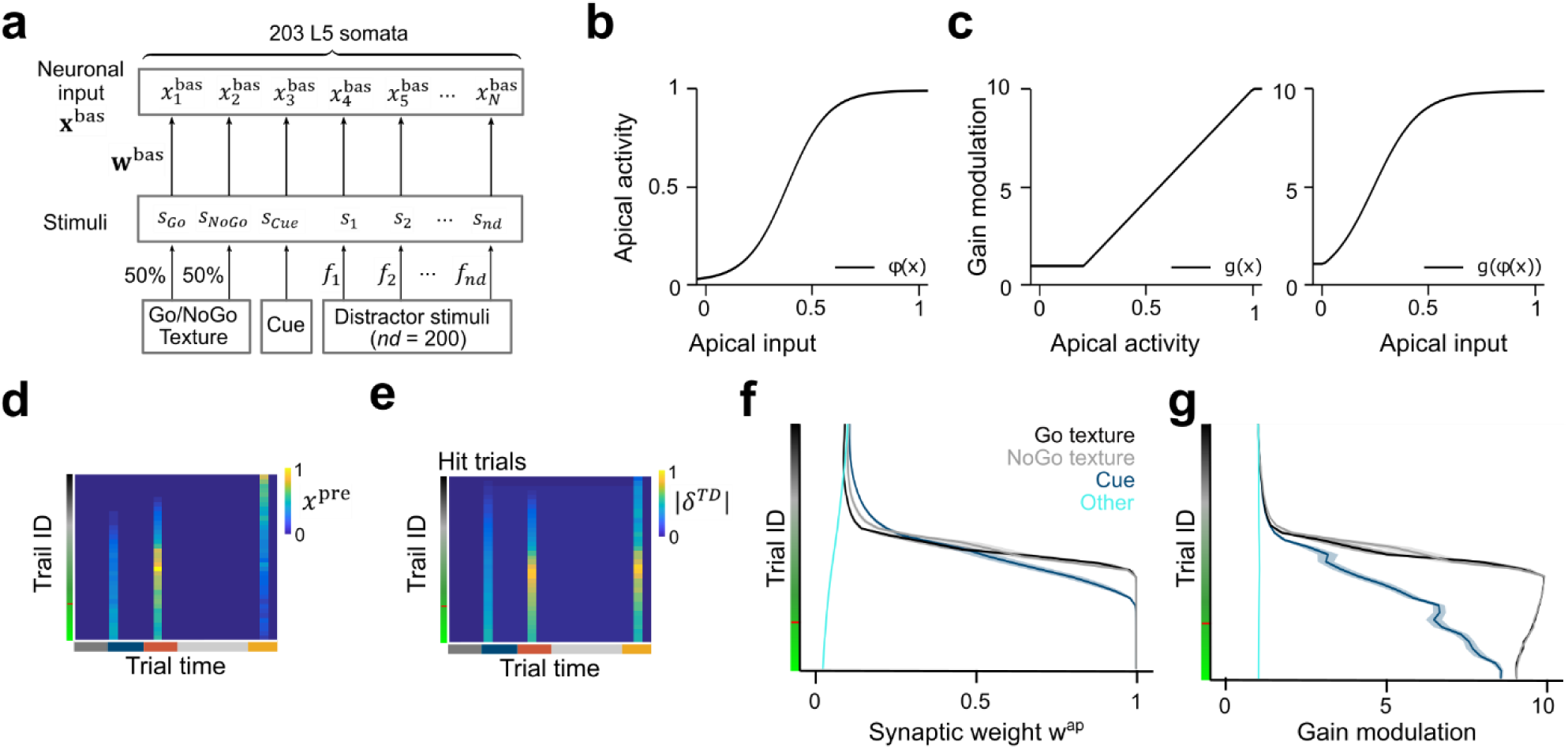
Temporal-difference learning model for dendritic salience proceesing. **a**, Detailed schematic of basal inputs to S1 L5 neurons in the pure-selectivity model. A total of N = 203 neurons was modelled. Besides Go, NoGo, and Cue stimuli other distracting stimuli were applied (number of distractors, *nd* = 200). **b**, Transfer function between apical input and apical dendritic activity. **c**, Dependence of gain modulation on apical activity and apical inputs. **d**, Evolution of the presynaptic input to sensory dendrites *x*^*pre*^, which represents an internal estimate of the conditioned salience, across learning in Hit trials. **e**, Evolution of the unsigned TD error across learning in Hit trials. **f,** Evolution of synaptic weights in cue, touch and outcome window across learning for Go, NoGo, Cue neurons and other distractor neurons. **g,** Evolution of the apical gain modulation across learning for Go, NoGo, Cue neurons and other distractor neurons.

**Supplementary Fig. 13.**
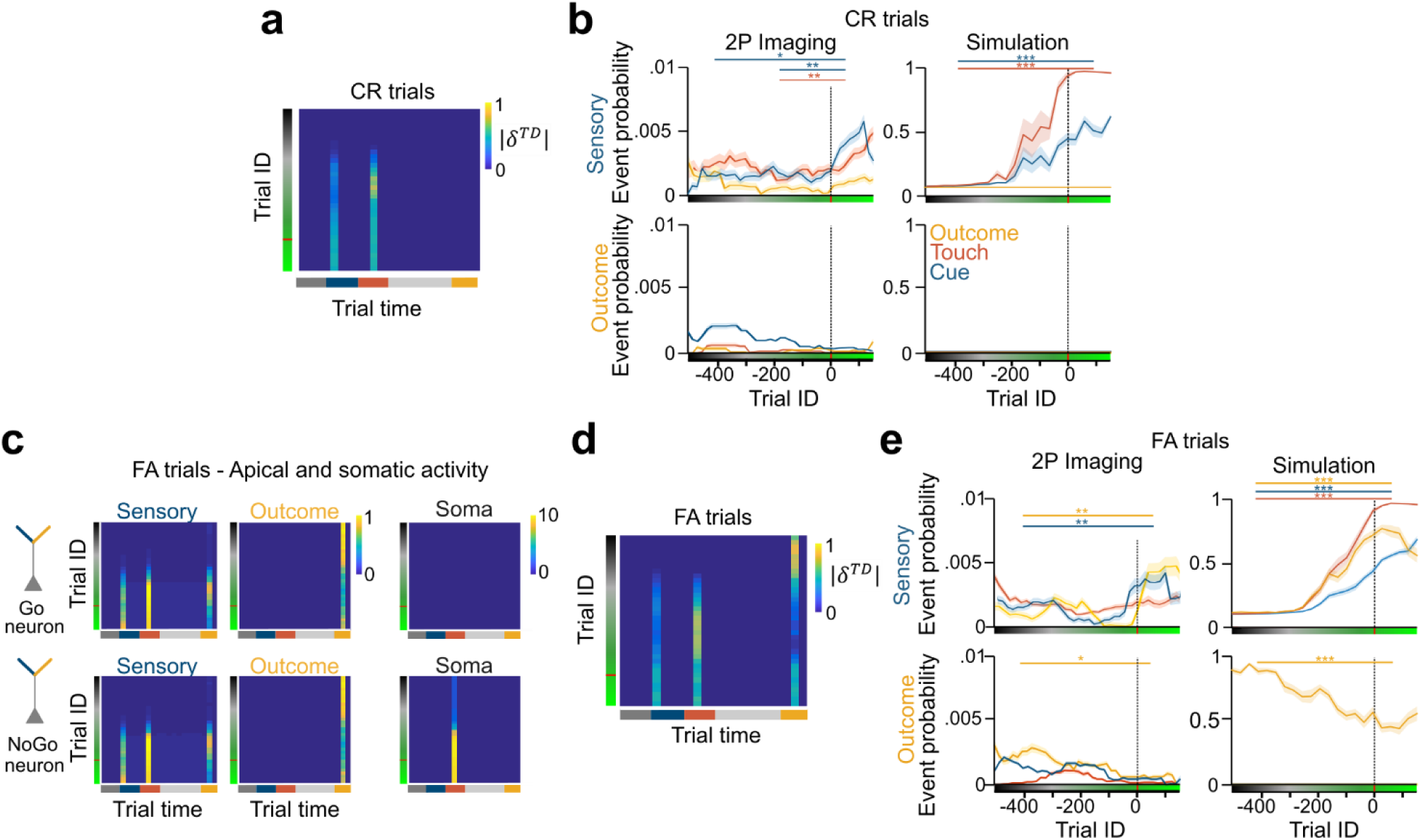
Computational model qualitatively reproduces 2P results of changes in event probability per trial window across learning. **A,** Unsigned TD error across learning in CR trials. **b,** Left: Event probability for event transient in CR trials in the Cue, Touch and Outcome window in sensory-driven and outcome-driven branches across learning based on 2P data and simulations (mean ± s.e.m., Sensory-driven branches, Cue window: naïve vs. expert, p= 0.016; learning vs. expert, p=0.004; Touch window: learning vs. expert, p=0.008). **c**, Apical activity of sensory and outcome branches as well as somata belonging to Go and NoGo neurons in FA trials. **d**, Unsigned TD error across learning in FA trials. **e**, Event probability in FA trials (Sensory-driven branches, Cue window: naïve vs. expert, p=0.005; Outcome window: naïve vs. expert, p=0.003; Outcome-driven branches, Outcome window: naïve vs. expert, p=0.04). For the simulations all comparisons indicated as significant had p<0.001. *p<0.05, **p<0.01, ***p<0.001

**Supplementary Fig. 14.**
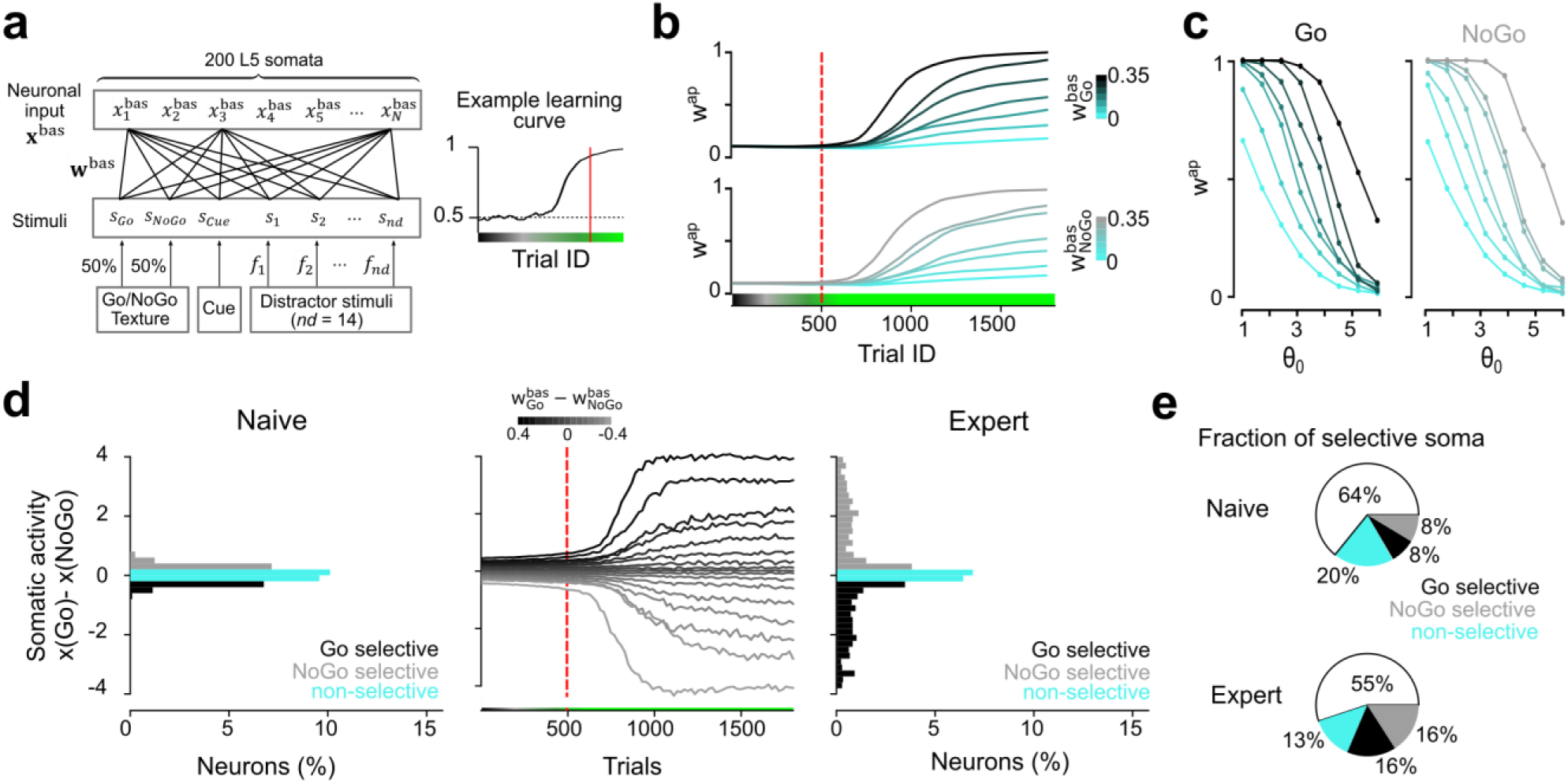
Computational model with mixed bottom-up selectivity. **a**, Schematic of the basal inputs’ connectivity in mixed-selectivity model. The basal synaptic weight matrix is randomly initialized and maps 3+nd inputs onto N L5 sensory pyramidal neurons (nd = 14; N = 200). **b**, Evolution of the synaptic weights onto sensory apical dendrites depending on the basal synaptic weight from the Go stimulus (top) or the NoGo stimulus (bottom). **c**, Strength of the synaptic weights onto sensory apical dendrites after 1800 trials dependent on the parameter θ_0_ (see Supplementary Notes) and on the basal synaptic weight from the Go stimulus (left) or the NoGo stimulus (right). **d**, Left: Histogram of the somatic selectivity of L5 pyramidal neurons towards Go/NoGo stimuli (quantified as the difference between the somatic responses to the Go and the NoGo stimulus) shown for the naïve state (before learning). Non-selective neurons are defined by somatic activity that neither favours Go nor NoGo texture (|x(Go)-x(NoGo)|<0.2) and neurons that were unresponsive to either stimulus were not shown. Inactive neurons were defined as neurons with a mean somatic response that was lesser than 0.1 during the stimulus window. Middle: Development of somatic response selectivity, depending on the basal selectivity towards Go/NoGo stimuli (quantified as the difference between the basal synaptic weights from the Go and NoGo stimulus). Right: Same as on the left but shown for the expert state (after 1800 trials). **e**, Fractions of neurons that are Go-selective, NoGo-selective, non-selective or inactive in the naïve and expert state.

**Supplementary Fig. 15.**
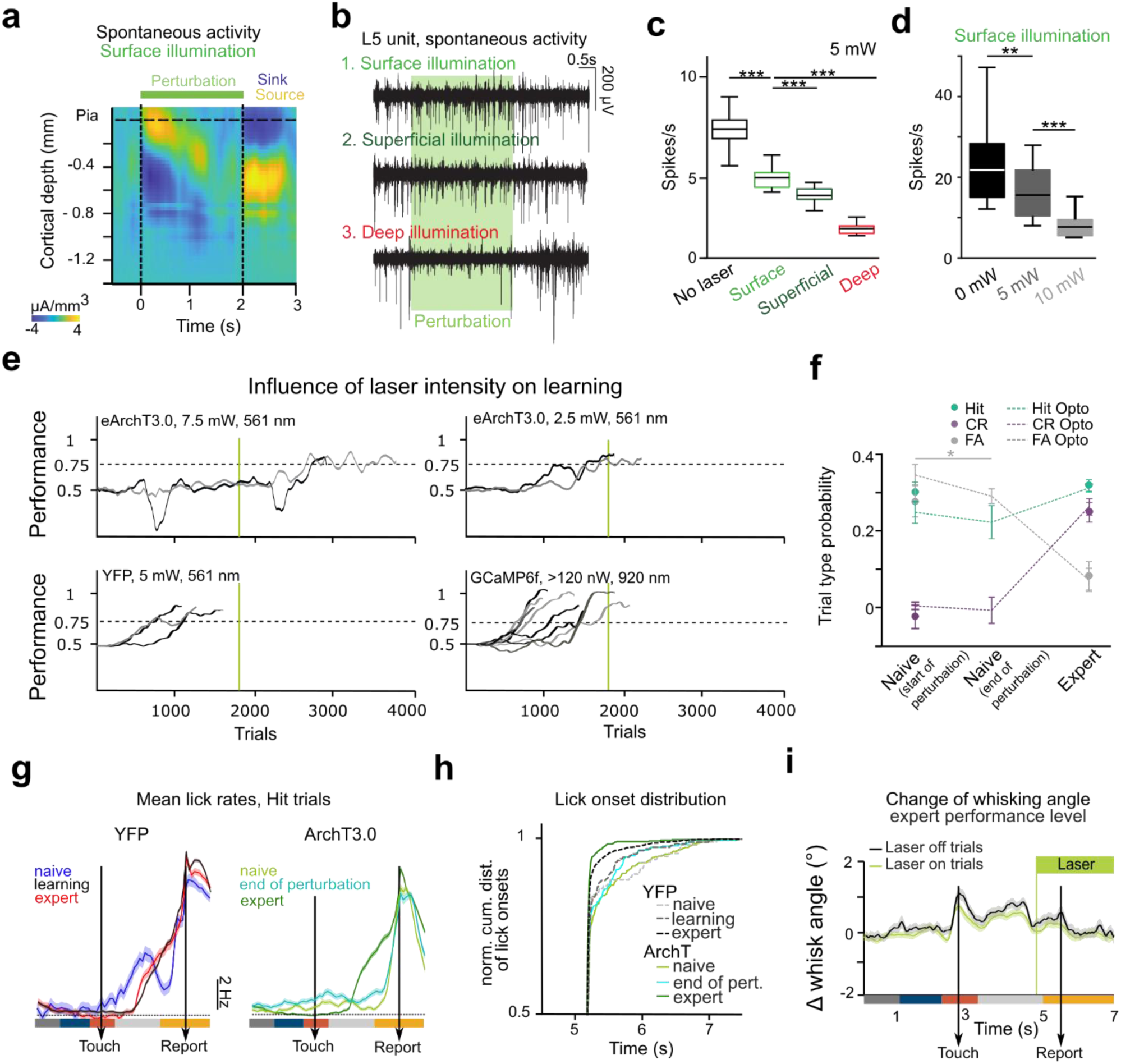
Behavioral and electrophysiological controls of optogenetic perturbation. **a**, Average current source density across S1 around optogenetic perturbation (isoflurane aesthesia, 3 sessions, 2 mice, 300 trials). **b**, Example of multi-unit activity from a single location showing illumination position dependent suppression of multi-unit activity. **c,** Firing rate of L5 units with and without optogenetic manipulation according to fiber position (line: median, box: 25^th^ and 75^th^ percentile, whiskers: 5^th^ and 95^th^ percentile; 5 mW, 17 units, 200 trials, 8 sessions, n = 4 mice, no laser vs. surface p=1e-12; surface vs. superficial p=8e-6; surface vs. deep p=0.9e-20; t-test). **d,** Firing rate of L5 units with and without optogenetic manipulation of the surface according to laser intensity (0, 5, or 10 mW, 17 units, 200 trials, 6 sessions, n = 4 mice; 0 mW vs. 5 mW p=0.0025; 5 mW vs. 10 mW p=0.0006, t-test). **e**, Learning curves of various control experiments with eArchT3.0-, eYFP- or GCaMP6f-expressing mice excited by 561-nm or 920-nm laser light with 2.5, 5.0, 7.5 or >120 mW light intensity (Green line: end of the laser perturbation after 1’800 trials). **f,** Probability for Hit, CR and FA trials in naïve and expert state of control mice (filled circles), compared to the respective probabilities in optogenetically stimulated mice in naïve state, at the end of perturbation, and in expert state (n = 8 and 5 mice for two-photon and optogenetic experiments;.Wilcoxon-ranksum test, p=0.031). **g**, Average lick rates during Hit trials in eYFP-expressing control mice (n = 3; 5 mW laser illumination at 561 nm; 321, 401 and 589 trials in the naïve, learning, and expert condition) and eArchT3.0-expressing mice (n = 5; 5 mW laser illumination, 1000 trials per condition) during and after perturbation (end of perturbation= last 200 perturbation trials, mean ± s.e.m). **h,** Comparison of normalized cumulative distribution of reward-triggering licking onsets in Hit trials in eArchT3.0- and eYFP-expressing mice. **i,** Average whisking angle in Hit trials in expert mice in laser on and laser off trials (468 and 428 trials; n = 3 mice). *p<0.05, **p<0.01, ***p<0.001

**Supplementary Video 1 | Example two-photon calcium imaging videos of trunk and apical tuft dendrites showing global and local events**. **a**, Example video showing a global event followed by a local event.*Left panel:* Motion-corrected calcium imaging movie of trunk cross-section in the lowest imaging plane (approx. 150 µm below the pia) recorded at 10 Hz. Scale bar is 50 µm. Trunk ROI is shown in blue. *Middle panel:* Motion-corrected calcium imaging movie of the most apical imaging plane (approx. 32 µm below the pia) recorded at 10 Hz. Scale bar is 50 µm. Example dendritic ROI is shown in red. *Right panel:* ΔF/F traces for the trunk (blue) and dendritic ROI (red). Scale bars are 2 s and 100%. **b**, Example video of a local event. **c**, Example video of a local event followed by a global event.

## Notes

### Competing Interest Statement

The authors have declared no competing interest.

### Summary of Updates

Revised manuscript including new experiments and analysis; addition of a computational model described in Supplementary Information file; 3 authors added.

