## Supplemental Notes for "Unsigned temporal difference errors in cortical L5 dendrites during learning"

### Supplementary Information

**Note 1. Error components of temporal difference learning map onto L5 apical tuft activity**

**Note 2. Computational model of gain modulation by unsigned TD errors in L5 dendritic tufts**

**Note 3. Apical dendrite activity amplifies task-relevant neurons**

**3.1 Convergence of the conditioned salience**

**3.2 Analyzing the definition of task relevance**

**3.3 Setting the plasticity threshold $\boldsymbol{\theta_i}$**

These notes expand on various aspects of the computational model. The first note recapitulates the mathematical theory of temporal difference (TD) learning and details how the TD error can be decomposed into interpretable components that characterize two types of salience. We map these salience terms onto the two classes of apical dendrite activity observed in our experiments. In the second note we provide the details of the computational model with key equations for the learning dynamics and the parameter settings for both the pure-selectivity and the mixed-selectivity model. In the final note we perform various in-depth analyses of the simulations regarding the convergence (or expert state) of the computational model, our formal definition of task relevance, and the selection of the plasticity threshold parameter.

#### Note 1. Error components in temporal difference learning map onto L5 apical tuft activity

To compare our experimental findings of learning-related dendrite dynamics to the TD error, we deem it useful to first provide a basic description of classic TD learning[1]. We will demonstrate how scrutinizing the expected reward $\hat{R}_t$ allows dissection of the TD error $\delta^{TD}$ into two components: an ***unconditioned error*** $\delta^{U}$ and a ***conditioned error*** $\delta^{C}$. We then relate these TD error components to salience terms and map them onto the dendritic signals observed in our two-photon calcium imaging data.

In discrete time formulation, the state-value function of an expected outcome is defined as

$$V_t(S_t) = \mathbb{E}[\sum_{k=0}^{\infty} \gamma^k R_{t+k+1} | S_t] \quad (1)$$

where $R_{t+1}$ is the reward outcome at time $t+1$, $S_t$ the state representation at time $t$, and $\gamma \epsilon [0,1]$ a discounting parameter for future rewards (in our model we assumed $\gamma = 1$, i.e., no discounting). Here, rewards $R$ can be either positive or negative in order to model both reward and punishment in our biological study. Typically, the state value is learned by an estimator $\hat{V}$ using the TD(0) error given by Sutton and Barto[1]:

$$\delta_t^{TD} = R_{t+1} + \gamma \hat{V}_{t+1}(S_{t+1}) - \hat{V}_t(S_t) \quad (2)$$

$$\Delta \mathbf{w}_{t+1}^V = \eta_V \, \delta_t^{TD} \frac{d}{d\mathbf{w}^V} \hat{V}_t(S_t) \quad (3)$$

Equation 3 implies that the weights $\mathbf{w}^V$, parametrizing the state-value estimator $\hat{V}$, are updated with the TD error $\delta^{TD}$ using a learning rate $\eta_V$. Note that we use the time index twice when writing $\hat{V}_t(S_t)$. This is redundant, however it is necessary in order to properly formulate the expected reward at time $t+1$ as the change in state-value estimate conditioned on state $S_t$ in the time step between $t$ and $t+1$:

$$\hat{R}_{t+1} = \hat{V}_t(S_t) - \gamma \hat{V}_{t+1}(S_t) \quad (4)$$

This definition is self-consistent for a state-value estimator $\hat{V}$ that consists of an explicit reward prediction at each time step

$$\hat{V}_t(S_t) = \sum_{k=0}^{\infty} \gamma^k \hat{R}_{t+k+1}(S_t) \quad (5)$$

This can be seen by pulling out the first summand and rearranging the equation:

$$\begin{aligned} \hat{R}_{t+1} &= \hat{V}_t(S_t) - \sum_{k=1}^{\infty} \gamma^k \hat{R}_{t+k+1}(S_t) \\ &= \hat{V}_t(S_t) - \gamma \sum_{k=0}^{\infty} \gamma^k \hat{R}_{t+k+2}(S_t) \\ &= \hat{V}_t(S_t) - \gamma \hat{V}_{t+1}(S_t) = -\Delta \hat{V}_{t+1}^R \end{aligned} \quad (6)$$

Here, we introduced $\Delta \hat{V}^R$ as the decrease in state-value estimation in time steps when a reward is expected. Assuming that such imminent reward predictions are modeled, changes in the state-value estimation from one time point $t$ to the next $t+1$ can be written as

$$\begin{aligned} \Delta \hat{V}_{t+1} &= \gamma \hat{V}_{t+1}(S_{t+1}) - \hat{V}_t(S_t) \\ &= \gamma \hat{V}_{t+1}(S_{t+1}) - \gamma \hat{V}_{t+1}(S_t) + \gamma \hat{V}_{t+1}(S_t) - \hat{V}_t(S_t) \\ &= \gamma \left( \hat{V}_{t+1}(S_{t+1}) - \hat{V}_{t+1}(S_t) \right) + (\gamma \hat{V}_{t+1}(S_t) - \hat{V}_t(S_t)) \\ &= \gamma \, \Delta \hat{V}_{t+1}^S + \Delta \hat{V}_{t+1}^R \\ &= \gamma \, \delta_{t+1}^C - \hat{R}_{t+1} \end{aligned} \quad (7)$$

Here, $\delta_{t+1}^C = \Delta \hat{V}_{t+1}^S = \hat{V}_{t+1}(S_{t+1}) - \hat{V}_{t+1}(S_t)$ represents the difference in state-value estimation from time $t$ to $t+1$ that results from updating the state from $S_t$ to $S_{t+1}$. While this double notation may seem redundant at first, the two terms reflect different connotations of this variable in different contexts: $\delta^C$ describes the conditioned error component of the TD error, whereas $\Delta \hat{V}^S$ describes the temporal changes of $\hat{V}$ that can be attributed to sensory stimuli.

Assuming that the estimator $\hat{V}$ is accurate (i.e., $\hat{V} = V$), $\delta_{t+1}^C$ is exclusively influenced by state changes that are informative for predicting rewards, i.e., conditioned stimuli (or actions) that change the estimated value. We therefore refer to $\delta_{t+1}^{C}$ as ***conditioned error***, which will turn out to be key for the detection of task-relevant stimuli (see below). By introducing an additional ***unconditioned error*** $\delta_{t+1}^{U} = R_{t+1} - \hat{R}_{t+1}$ we can dissect the TD error $\delta_t^{TD}$ into two error components:

$$
\begin{aligned}
\delta_t^{TD} &= R_{t+1} + \gamma \hat{V}_{t+1}(S_{t+1}) - \hat{V}_t(S_t) \\
&= R_{t+1} + \Delta \hat{V}_{t+1} \\
&= R_{t+1} + \gamma\, \Delta \hat{V}_{t+1}^{S} + \Delta \hat{V}_{t+1}^{R} \\
&= R_{t+1} - \hat{R}_{t+1} + \gamma\, \delta_{t+1}^{C} \\
&= \delta_{t+1}^{U} + \gamma\, \delta_{t+1}^{C}
\end{aligned}
\tag{8}
$$

We relate these TD error components to the two functional classes of learning-associated dendritic branch dynamics that we have revealed in our study. First, any unexpected event of high motivational relevance (reward or punishment) is associated with ***unconditioned salience*** $s_t^{U}$, which we define as the absolute value of the unconditioned error $\delta_t^{U}$

$$
s_t^{U} = |\delta_t^{U}| = |R_t - \hat{R}_t| \tag{9}
$$

Second, any sensory information that bears predictive power for future relevant events over time will evoke ***conditioned salience*** $s_t^{C}$, defined as the unsigned change of the current state-value estimate**:**

$$
s_t^{C} = |\Delta \hat{V}_t| = |\gamma\, \hat{V}_t(S_t) - \hat{V}_{t-1}(S_{t-1})| = \left|\gamma\, \delta_t^{C} - \hat{R}_t\right| \tag{10}
$$

This term includes the conditioned error $\delta_t^{C}$, which quantifies changes in state-value estimate due to sensory information, and the reset $(-\hat{R}_t)$ when the outcome is expected. We interpret the two functional classes of learning-associated dendritic branch dynamics as reflecting excitatory responses that represent these two types of salience terms, which are based on unsigned TD error components. Specifically, we associate the responses in outcome branches with the unconditioned salience $s_t^{U}$ and the activity in sensory branches with the conditioned salience $s_t^{C}$.

### Note 2. Computational model of gain modulation by unsigned TD errors in L5 dendritic tufts

Here we explain the details of our computational model (see also Fig. 3b and Supplementary Figs. 12-14). We modeled a population of 203 rate-based L5 neurons. They receive basal inputs directly onto their soma and apical inputs onto two distinct tuft branches of the apical dendrite. The simulations are centered around modulating the somatic activities $\mathbf{x}^{\mathrm{som}}$ of pyramidal neurons, driven by fixed basal synapses $\mathbf{w}^{\mathrm{bas}}$, with a multiplicative gain imposed via the apical activity $\mathbf{x}^{\mathrm{ap}}$. The apical activity is modeled as the sum of activities of the two dendritic compartments $\mathbf{d}^{\mathrm{sen}}$ and $\mathbf{d}^{\mathrm{out}}$, representing sensory and outcome branch types and each receiving a distinct type of input. Each behavioral trial is modeled with time steps from $t = 1 \ldots t_R$ with $t_R = 18$ as the last time step of the trial, during which the outcome $R$ is realized according to the trial type (Hit: reward $R = 1$; CR and Miss: no reward $R = 0$; FA: light punishment $R = -0.5$).

***Basal inputs.*** Basal inputs $\mathbf{x}^{\text{bas}}$ convey information about various sensory stimuli to the somata of the model neuron population. Sensory stimuli are modeled as binary variables that are either activated once during a given trial or absent from the trial. These stimuli comprise three task-relevant sensory variables (go-texture, no-go-texture, tone-cue) as well as various distractor stimuli. The tone-cue is active in the first time step of each trial and the texture stimuli are randomly alternated and always confined to the texture time step. The distractor stimuli were randomly sampled from individual Bernoulli distributions with different linearly spaced activation probabilities between 0 and 1. The activation timings of the distractor stimuli were random and uniformly drawn across the trial. In the **pure-selectivity model**, each of the three task-relevant stimuli and the 200 distractor stimuli were connected each to the soma of one of the 203 L5 pyramidal neurons (Supplementary Fig. 12a); in other words, the basal synaptic weight matrix was an identity matrix.

In the **mixed-selectivity model**, the basal stimuli were mapped to the L5 neurons in a distributed manner, with the basal weights probabilistically drawn (Supplementary Fig. 14a) according to the distribution

$$\mathrm{p}\left(\mathrm{w}_{\mathrm{ij}}^{\mathrm{bas}}\right) = \begin{cases} \frac{1}{\alpha}(\mathrm{w}_{\mathrm{ij}}^{\mathrm{bas}})^{1/\alpha-1} & \text{if } 0 \leq w_{ij}^{\mathrm{bas}} \leq 1 \\ 0 & \text{otherwise} \end{cases} \tag{11}$$

with $\alpha = 6$. After sampling, the weights to each neuron $i$ were normalized to sum up to 1 and remained fixed throughout the entire simulation. In the mixed-selectivity model, only 14 distractor stimuli were used.

***Apical dendritic inputs***. Each neuron is modeled with two apical dendritic compartments $d_i^{\text{sen}}$ and $d_i^{\text{out}}$, prototypical for the identified functional branch types. All outcome branches across the modelled population uniformly received the same input

$$d_{i,t}^{\text{out}} = \varphi(s_t^U) = \varphi\left(\left|R_t - \hat{R}_t\right|\right) \quad \forall\, i \tag{12}$$

where $s_t^U$ is the unconditioned salience and $\varphi$ is a sigmoidal transfer function (Supplementary Fig. 12b). The activity of the **sensory branches** is modeled as a neuron-specific conditioned salience signal

$$d_{i,t}^{\text{sen}} = \varphi\left(w_{i,t}^{\text{ap}}\, x_t^{\text{pre}}\right) \tag{13}$$

Here, the temporally fluctuating input $x_t^{\text{pre}}$ is modeled to approximate conditioned salience, i.e., $x_t^{\text{pre}} \approx \left|\Delta \hat{V}_t\right|$, while the neuron-specific weights $w_{i,t}^{\text{ap}}$ are adjusted during learning to steer $x_t^{\text{pre}}$ predominantly towards task-relevant neurons (see note 3.2, equation 34). Specifically, the presynaptic activity $x_t^{\text{pre}}$ is modeled as

$$x_t^{\text{pre}} = \gamma\, \hat{s}_t^{\text{stim}} + \left|\hat{R}_t\right| \tag{14}$$

where $\hat{s}_t^{\text{stim}}$ represents the salience induced by sensory stimuli. Hence, unsigned changes $\Delta \hat{V}_t$ are modeled in $x^{\text{pre}}$ by separately predicting components that arise from sensory information ($\hat{s}^{\text{stim}}$) and components that arise from performing actions (retrieving $\hat{R}$). $\hat{s}^{\text{stim}}$ is modeled as a time-dependent estimator for $\mathbb{E}\left[\left|\delta_t^C\right|\right]$, the expected unsigned fluctuation in $\hat{V}$ attributed to sensory information in the somatic activities $\mathbf{x}^{\text{som}}$ (see equation 7) during each time step. This estimator is updated according to

$$\Delta\hat{s}_t^{\text{stim}} = \eta_a\left(\left|\delta_t^C\right| - \hat{s}_t^{\text{stim}}\right) \tag{15}$$

where $\eta_a$ denotes the learning rate of the afferent input. The weights $w_{i,t}^{\text{ap}}$ were updated individually for each neuron $i$ with a local learning rule

$$\Delta w_{i,t}^{\text{ap}} = \eta_w\, \kappa\left(w_{i,t}^{\text{ap}}\right)\left(x_{i,t}^{\text{som}} - \theta_i\right) x_t^{\text{pre}} \tag{16}$$

$$\kappa\left(w_{i,t}^{\text{ap}}\right) = \left(w_{i,t}^{\text{ap}} - w_{min}\right)\left(w_{max} - w_{i,t}^{\text{ap}}\right) \tag{17}$$

where $\eta_w$ denotes the apical learning rate and $\theta_i$ the fixed plasticity threshold (see note 3.3). The apical weights are adapted online and stayed bound within $[w_{\min}, w_{\max}]$ due to the term $\kappa\left(w_{i,t}^{\text{ap}}\right)$. With these definitions, the sensory branch excitation $w_{i,t}^{\text{ap}} x_t^{\text{pre}}$ can be shown to converge towards the conditioned salience $s_t^C$, if a neuron $i$ is task-relevant and towards zero otherwise (see note 3.1 below; equation 33).

The overall apical tuft activity is modeled as the sum of the activity in the outcome and sensory branch for each neuron

$$\mathbf{x}^{\text{ap}} = \mathbf{d}^{\text{out}} + \mathbf{d}^{\text{sen}} \tag{18}$$

***Gain modulation of somatic activity*.** As core feature in our model, the basally driven somatic activities $\mathbf{x}^{\text{som}}$ are gain modulated via the apical dendritic activity, i.e.,

$$x_{i,t}^{\text{som}} = g_{i,t}\left(x_{i,t}^{\text{bas}} + b\right) \tag{19}$$

where $x_{i,t}^{\text{bas}}$ are the basal inputs, as given above for either the pure-selectivity or the mixed-selectivity model, and $b$ is a constant background activation. The apical gain $g_i$ is given by

$$g_{i,t} = \begin{cases} 1 + \left(g_{\max} - 1\right) \cdot x_{i,t}^{\text{ap}} & \text{if } 0 \le x_{i,t}^{\text{ap}} \le 1 \\ g_{\max} & \text{if } x_{i,t}^{\text{ap}} > 1 \end{cases} \tag{20}$$

Here, we chose $g_{\max} = 10$. The history of the somatic output $\mathbf{x}^{\text{som}}$ over all timesteps within the current trial is integrated into an eligibility trace $\mathbf{z}$ according to

$$\mathbf{z}_t = \mathbf{z}_{t-1} + \Delta\mathbf{z} = \mathbf{z}_{t-1} + \mathbf{x}_t^{\text{som}}, \tag{21}$$

which is reset to zero between trials.

***Reinforcement learning*.** Our model is based on TD learning of the state-value function $V$ with an estimator

$$\begin{aligned} \hat{V}_{\text{t}} &= \gamma^{-1}\,\hat{V}_{\text{t}-1} + \gamma^{-1}\Delta\hat{V}_t \\ &= \gamma^{-1}\,\hat{V}_{\text{t}-1} + \delta_t^C - \gamma^{-1}\hat{R}_t \\ &= \textstyle\sum_{t'=1}^{t}\left(\gamma^{t'-t}\,\delta_{t'}^C - \gamma^{t'-t-1}\hat{R}_{t'}\right) \end{aligned} \tag{22}$$

which is consistent with equation 7. Instead of estimating the value $\hat{V}_{\text{t}}$ directly based on a state representation, our model integrates the estimation of the changes of the value function $\Delta\hat{V}_t$ over time. The changes in value expectation are either based on sensory information ($\delta_t^C$) or on imminent expected reward ($\hat{R}_t$). The estimator $\delta_t^C$ is implemented as a perceptron network using the activity pattern of sensory pyramidal neuron $\mathbf{x}^{\text{som}}$ as input:

$$\delta_t^C = \mathbf{w}_t^V \cdot \mathbf{x}_t^{\text{som}} \tag{23}$$

The network's synaptic weights $\mathbf{w}^V$ are updated with a learning rule based on the TD-delta decomposition (equation 8):

$$\begin{aligned}\Delta\mathbf{w}_t^V &= \eta_V((R_t - \hat{R}_t)(\mathbf{x}_t^{\text{som}} - b) \odot \mathbf{z}_{t-1} + k^V\gamma\delta_t^C\mathbf{z}_{t-1} + k^V\delta_t^{reset}\mathbf{z}_{t-1}) \\ &= \eta_V((R_t - \hat{R}_t)(\mathbf{x}_t^{\text{som}} - b) + k^V\gamma\delta_t^C + k^V\delta_t^{reset}) \odot \mathbf{z}_{t-1} \\ &= \eta_V((R_t - \hat{R}_t)(\mathbf{x}_t^{\text{som}} - b) + k^V\gamma\delta_t^C + k^V\delta_t^{reset}) \odot \frac{d}{d\mathbf{w}^V}\textstyle\sum_{t'=1}^{t-1}\delta_{t'}^C \\ &= \eta_V((R_t - \hat{R}_t)(\mathbf{x}_t^{\text{som}} - b) + k^V\gamma\delta_t^C + k^V\delta_t^{reset}) \odot \frac{d}{d\mathbf{w}^V}\hat{V}_{t-1}\end{aligned} \tag{24}$$

where $\eta_V$ is the learning rate, $k^V = b \cdot g_{max} = 0.1$ is a constant, $\hat{R}$ and $\delta_t^{reset}$ are defined below. This approach resembles traditional TD learning (see equation 3) with the exception that the conditioned and the unconditioned components of the TD error are treated separately. The imminent reward prediction $\hat{R}$ is derived from the state-value estimate by resetting $\hat{V}$ to zero in the next time step, if the lick action is performed. This leads to

$$\hat{R}_{t+1} = \begin{cases} \hat{V}_t & \text{if } t = t_R - 1 \text{ and } a_{t_R-1} = a_{go} \\ 0 & \text{otherwise} \end{cases} \tag{25}$$

which is in line with the self-consistency equation for $\hat{R}$ (equation 6). The additional update $\delta_t^{reset}$ is applied to reduce any remaining error in the state-value estimation at the end of each trial, which is equivalent to the TD error $\delta_t^{TD} = \hat{R}_{t+1} + \gamma\hat{V}_{t+1} - \hat{V}_t$ resulting from resetting the estimate $\hat{V}_{t+1}$ to zero after a trial is finished.

$$\delta_t^{reset} = \begin{cases} -\hat{V}_t & \text{if } t = t_R \\ 0 & \text{otherwise} \end{cases} \tag{26}$$

The **action selection network** consists of a perceptron with a sigmoid transfer function $\phi(x) = 1/(1+e^{-x})$, which integrates sensory inputs $\mathbf{x}_\text{t}^{\text{som}}$ across time to derive the go-action (lick) probability

$$\pi(\text{a}_{\text{go}}|\mathbf{x}_{1:\text{t}}^{\text{som}}) = \phi(\textstyle\sum_{t'=1}^{t}\mathbf{w}^\pi \cdot \mathbf{x}_{t'}^{\text{som}}) = 1 - \pi(a_{\text{nogo}}|\mathbf{x}_{1:\text{t}}^{\text{som}}) \tag{27}$$

This policy network has synapses $\boldsymbol{w}^\pi$ that are trained according to the policy gradient[1] weighted by the presynaptic reactivation $\mathbf{x}^{\text{som}}$

$$\begin{aligned}\Delta\mathbf{w}^\pi &= -\eta_\pi R_t \boldsymbol{x}_t^{\text{som}} \frac{d}{d\mathbf{w}^\pi}\log\big(\pi(a_{t-1}|\mathbf{x}_{1:\text{t}-1}^{\text{som}})\big) \\ &= -\eta_\pi R_t(1 - \pi(\text{a}_{\text{t}-1}|\mathbf{x}_{1:\text{t}-1}^{\text{som}}))\boldsymbol{x}_{\boldsymbol{t}}^{\text{som}} \odot \mathbf{z}_{t-1}\end{aligned} \tag{28}$$

where $\eta_\pi$ denotes the policy learning rate and $\text{a}_{\text{t}-1}$ the action ($\text{a}_{\text{go}}$ or $a_{\text{nogo}}$) performed in the previous time step.

***Apical inhibition.*** The replicate the experimental inhibition of outcome-related activity using optogenetics, we applied the equivalent perturbation in the computational model during the last time step of the simulated trials, i.e., the apical activities were set to zero for the time step at which the reward was delivered ($\mathbf{x}_{t_R}^{\mathrm{ap}} = 0$). This inhibition was lifted after 1800 trials and the simulation continued for 2200 further trials.

***Simulation parameters***. The apical weights for sensory apical inputs $\mathbf{w}^{\mathrm{ap}}$ were initialized according to a uniform distribution $\mathcal{U}(0.05, 0.15)$. The weights of the policy $\mathrm{w}_{ij}^{\pi}$ and of the state-value estimator $\mathrm{w}_{ij}^{V}$ were all initialized to zero. The numerical values for the learning rates were all set equal ($\eta_V = \eta_\pi = \eta_w = \eta_\mathrm{a}$ ) and chosen such that the simulations matched the number of trials that the mice needed to reach expert performance in the biological experiments. The values of all parameters necessary for the computational simulations are summarized in Supplementary Table 1.

| Parameter description | Symbol | Value |
|---|---|---|
| Hit reward | $R_{\mathrm{Hit}}$ | 1 |
| CR and Miss outcome | $R_{\mathrm{CR}}, R_{\mathrm{Miss}}$ | 0 |
| False alarm punishment | $R_{\mathrm{FA}}$ | -0.5 |
| State-value learning rate | $\eta_V$ | 0.016 |
| Policy learning rate | $\eta_\pi$ | 0.016 |
| Apical synapse weight learning rate | $\eta_w$ | 0.016 |
| Apical afferent input learning rate | $\eta_\mathrm{a}$ | 0.016 |
| Minimum value of $w^{\mathrm{ap}}$ | $w_{\mathrm{min}}$ | 0 |
| Maximum value of $w^{\mathrm{ap}}$ | $w_{\mathrm{max}}$ | 1 |
| Discount factor | $\gamma$ | 1 |
| Basal activity | $b$ | 0.01 |
| Maximal gain | $g_{\mathrm{max}}$ | 10 |
| Apical weight plasticity threshold | $\theta_i$ | 0.1 |

**Supplementary Table 1**: Model initialization parameters and the values chosen in the pure-selectivity model. In the mixed-selectivity model all parameters were the same except for the apical weight plasticity threshold $\theta_i$, see note 3.3.

### Note 3. Apical dendrite activity amplifies task-relevant neurons

In this last note we analyse how our model converges to amplify the gain of task-relevant neurons: First, we explain how the conditioned salience signal is guided onto the sensory branches of task-relevant neurons by adjusting the apical synaptic weights $\mathbf{w}^{\mathrm{ap}}$. Second, we analyze the definition of neuron-specific task relevance that dictates the dynamics of the apical synapse plasticity. Lastly, we discuss the range of values for the parameter $\theta_i$, which determines the task-relevance threshold for the convergence of the apical synaptic weights.

#### 3.1 Convergence of the conditioned salience

Given a fixed state-value estimator $\hat{V}$ and policy $\pi$, one can derive the convergence state $x_t^{\mathrm{pre}*}$ for the presynaptic input of the conditioned salience model. After convergence we have

$$0 = \Delta\hat{s}_t^{\mathrm{stim}} = \eta_a\left(\left|\delta_t^C\right| - \hat{s}_t^{\mathrm{stim}}\right) \tag{29}$$

$$x_t^{\mathrm{pre}*} = \gamma\hat{s}_t^{\mathrm{stim}*} + \left|\hat{R}_t\right| = \gamma\left|\delta_t^C\right| + \left|\hat{R}_t\right| \approx \left|\Delta\hat{V}_t\right| = \mathrm{s}_t^C \tag{30}$$

with asterisks indicating converged values. The third step in equation 30 is accurate for our task and not just an approximation, because the model does not receive any new incoming sensory stimuli in the time step when the outcome is received, i.e., $\left|\delta_t^C\right|$ and $\left|\hat{R}_t\right|$ are never simultaneously non-zero. $x_t^{\mathrm{pre}*}$ can now be used to update the synaptic weights during learning:

$$\begin{aligned}
\mathbb{E}\left[\Delta w_{i.t}^{\mathrm{ap}}\right] &= \eta_w \cdot \mathbb{E}\left[\kappa\left(w_i^{\mathrm{ap}*}\right) \cdot x_t^{\mathrm{pre}*} \cdot \left(x_{i,t}^{\mathrm{som}} - \theta_i\right)\right] \\
&= \eta_w \cdot \kappa\left(w_i^{\mathrm{ap}*}\right) \cdot \mathbb{E}\left[s_t^C \cdot \left(x_{i,t}^{\mathrm{som}} - \theta_i\right)\right] \\
&= \eta_w \cdot \kappa\left(w_i^{\mathrm{ap}*}\right) \cdot \left(\mathrm{Cov}\left(s_t^C, x_{i,t}^{\mathrm{som}}\right) - \mathbb{E}\left[s_t^C\right]\left(\theta_i - \mathbb{E}\left[x_{i,t}^{\mathrm{som}}\right]\right)\right) \\
&\propto \underbrace{\left(\frac{\mathrm{Cov}\left(s_t^C, x_{i,t}^{\mathrm{som}}\right)}{\mathbb{E}\left[s_t^C\right]\mathbb{E}\left[x_{i,t}^{\mathrm{som}}\right]}\right)}_{\rho_i} - \underbrace{\left(\frac{\theta_i}{\mathbb{E}\left[x_{i,t}^{\mathrm{som}}\right]} - 1\right)}_{\rho_i^{th}}
\end{aligned} \tag{31}$$

This result enables us to derive the convergence of the apical top-down synaptic weights converge with learning, in terms of the ***task relevance*** $\rho_i$ of neuron $i$ (see note 3.2 below):

$$w_i^{\mathrm{ap}*} = \begin{cases} w_{\max} = 1 \ ; & \text{if } \rho_i > \rho_i^{th} \\ w_{\min} = 0 \ ; & \text{if } \rho_i < \rho_i^{th} \end{cases} \tag{32}$$

This result means that $x_t^{\mathrm{pre}}$ will preferably influence the apical dendrites of neurons, for which $\rho_i$ exceeds the threshold $\rho_i^{th}$. Combining equations 30 and 32, we find that the activities of sensory dendritic branches converge towards

$$d_i^{\mathrm{sen}*} = \varphi(w_i^{\mathrm{ap}*}\, x_t^{\mathrm{pre}*}) = \begin{cases} \varphi(\mathrm{s}_t^C) & ; \text{if } \rho_i > \rho_i^{th} \\ \varphi(0) & ; \text{if } \rho_i < \rho_i^{th} \end{cases} \tag{33}$$

In summary, if the task relevance $\rho_i$, which is determined by the covariance between $s_t^C$ and $x_i^{\mathrm{som}}$, is greater than a threshold $\rho_i^{th}$, the apical gain $g_i$ of this neuron $i$ will be strongly upmodulated by $x^{\mathrm{pre}}$. If the task relevance of a neuron is low, the gain will be downmodulated. Additionally, $x^{\mathrm{pre}}$ learns to predict time points, at which useful information is obtained. The reason is that at time points when stimuli are perceived that are informative for outcome prediction, the state-value prediction $\hat{V}$ will be affected. Here it is worth noting that combining a local plasticity rule with a pre-synaptic activity representing the temporal changes of $\hat{V}$ (encoding the temporal credit) provides a framework for spatial credit assignment of task relevance.

#### 3.2 Analyzing the definition of task relevance

In the above derivation, we defined the perceived task relevance for a neuron $i$ as

$$\rho_i = \mathrm{Cov}\left(\frac{|\Delta\hat{V}_t|}{\mathbb{E}[|\Delta\hat{V}_t|]}, \frac{x_{i,t}^{\mathrm{som}}}{\mathbb{E}[x_{i,t}^{\mathrm{som}}]}\right) = \mathrm{Cov}\left(\frac{s_t^C}{\mathbb{E}[s_t^C]}, \frac{x_{i,t}^{\mathrm{som}}}{\mathbb{E}[x_{i,t}^{\mathrm{som}}]}\right) \tag{34}$$

The task relevance $\rho_i$ can also be described as the covariance between the unsigned $\hat{V}$ fluctuations ($s_t^C = |\Delta\hat{V}_t|$) and the somatic activity of a neuron $i$, normalized by the average fluctuation level and the average somatic activity of the neuron $i$. In this section, we will attempt to provide some useful intuition on how to interpret this mathematical object. Before that we first introduce the notion of $\Delta\hat{V}_i^S$ as an estimate of the expected information gained by the system from the somatic activity of the neuron $i$. We can decompose $\Delta\hat{V}_t^S$ as follows:

$$\Delta\hat{V}_t^S = \delta_t^C = \sum_{i=1}^N \mathrm{w}_{i,t}^V \cdot x_{i,t}^{\mathrm{som}} = \sum_{i=1}^N \underbrace{\frac{\partial \delta_t^C}{\partial x_{i,t}^{\mathrm{som}}} \cdot x_{i,t}^{\mathrm{som}}}_{\Delta\hat{V}_{i,t}^S} = \sum_{i=1}^N \Delta\hat{V}_{i,t}^S \tag{35}$$

Here we introduced $\Delta\hat{V}_i^S$ as the component of the conditioned error that is contributed by neuron $i$. From now on moving forward, we will stop specifying the $t$-indices in every calculation step and set $\gamma = 1$ to simplify the notation. With the definition of $\Delta\hat{V}_i^S$ in equation 35, we can rewrite equation 11 as

$$s^C = |\Delta\hat{V}| = \left|\Delta\hat{V}^S - \hat{R}\right| = \left|\sum_{i=1}^N \Delta\hat{V}_i^S - \hat{R}\right| \tag{36}$$

Hence, at each time step one can compute the portion $\Delta\hat{V}_i^S$ of the state-value estimation update $\Delta\hat{V}$ that can be attributed to neuron $i$. While intuitively $\Delta\hat{V}_i^S$ might seem a promising basis to define the task relevance of a neuron, it is not the most adequate. Yet, we will use it as a starting point to explain why $\mathrm{Cov}(|\Delta\hat{V}|, x_i^{\mathrm{som}})$ is actually best suited for this purpose. For the analysis, we will assume $\mathrm{Cov}(|\Delta\hat{V}|, x_i^{\mathrm{som}}) = \mathrm{Cov}(|\Delta\hat{V}^S|, x_i^{\mathrm{som}})$, which is justified as long as stimuli do not co-occur with the timing of the outcome (when $\hat{R}_t \neq 0$).

The covariance $\mathrm{Cov}(|\Delta\hat{V}|, x_i^{\mathrm{som}})$ between a neuron's somatic activity and changes in the general prediction of future relevant outcomes (as conveyed to the apical tuft) is central to our definition of task relevance $\rho_i$. It is therefore useful to analyze this covariance term in more detail to provide some intuition for what it represents. Before considering the covariance with the absolute value ($|\Delta\hat{V}|$), it is simpler to look at the signed case, where we can make use of the distributivity of covariance:

$$\begin{aligned} \mathrm{Cov}(\Delta\hat{V}, x_i^{\mathrm{som}}) &= \mathrm{Cov}\left(\sum_j \Delta\hat{V}_j^S, x_i^{\mathrm{som}}\right) \\ &= \sum_j \mathrm{Cov}(\Delta\hat{V}_j^S, x_i^{\mathrm{som}}) \\ &= \sum_j \mathrm{Cov}\left(\frac{\partial \delta^C}{\partial x_j^{\mathrm{som}}} x_j^{\mathrm{som}}, x_i^{\mathrm{som}}\right) \\ &= \sum_j \mathbb{E}\left[\frac{\partial \delta^C}{\partial x_j^{\mathrm{som}}}\right] \mathrm{Cov}(x_j^{\mathrm{som}}, x_i^{\mathrm{som}}) \end{aligned} \tag{37}$$

For the last step, we need to assume that $\frac{\partial \delta^C}{\partial x_j^{\mathrm{som}}}$ and $x_i^{\mathrm{som}}$ are independent

$$0 = \mathrm{Cov}\left(\frac{\partial \delta^C}{\partial x_j^{\mathrm{som}}}, x_i^{\mathrm{som}}\right) \tag{38}$$

In our implementation, this is trivial since $\partial\delta^C / \partial x_i^{\mathrm{som}} = \mathrm{w}_i^V$. Equation 38 tells us that a neuron $i$ might become task relevant even if its own direct contributing factors $\Delta\hat{V}_i^S$ is small (as estimated by the state-value estimation model) as long as it covaries with one or several other neurons that strongly and directly contribute to the state-value estimation. Hence, one can remark that $\mathrm{Cov}(\Delta\hat{V}, x_i^{\mathrm{som}})$ makes a better definition of task relevance than $\mathrm{Cov}(\Delta\hat{V}_i^S, x_i^{\mathrm{som}})$, because the model $\hat{V}$ can contain redundancies or lack information that is contained in the full $\Delta\hat{V}$. For completeness, we can also look at the idealized situation, where $x_i^{\mathrm{som}}$ and $x_j^{\mathrm{som}}$ are independent for all $i \neq j$

$$\mathrm{Cov}\left(\Delta\hat{V}, x_i^{\mathrm{som}}\right) = \mathrm{Cov}\left(\Delta\hat{V}_i^S, x_i^{\mathrm{som}}\right) = \mathbb{E}\left[\frac{\partial\delta^C}{\partial x_i^{\mathrm{som}}}\right] \cdot \mathrm{Var}(x_i^{\mathrm{som}}) \tag{39}$$

Unfortunately, we cannot perform the equivalent derivations with the covariance of unsigned sums due to the lack of the distributive property. Hence, we must be content with looking at the intuitions that we gained and apply them to the unsigned case. This means that $\mathrm{Cov}(|\Delta\hat{V}|, x_i^{\mathrm{som}})$ detects task-relevant neurons, even if the state-value estimator $\hat{V}$ does not explicitly use their activities, and it ignores neurons that have a high $|\partial\delta^C/\partial x_j^{\mathrm{som}}|$ if they do not contribute to an actual change of estimate $|\Delta\hat{V}|$. This can be illustrated with some practical examples. By assuming that the state-value estimate $\hat{V}_1$ depends only on one neuron $k$, meaning that $\partial\delta^C/\partial x_j^{\mathrm{som}}| = 0\ \forall j \neq k,$ we find

$$\mathrm{Cov}\left(|\Delta\hat{V}_1|, x_i^{\mathrm{som}}\right) = \mathrm{Cov}\left(\left|\sum_j \frac{\partial\delta^C}{\partial x_j^{\mathrm{som}}} x_j^{som}\right|, x_i^{\mathrm{som}}\right) = \mathrm{Cov}\left(\Delta\hat{V}_{\mathrm{k}}^S, x_i^{\mathrm{som}}\right) \tag{40}$$

Using that $|\partial\delta^C/\partial x_k^{\mathrm{som}}|$ and $x_k^{\mathrm{som}}$ are independent and that $x_i^{\mathrm{som}} \geq 0$ for all $i$, we obtain

$$\mathrm{Cov}\left(|\Delta\hat{V}_1|, x_i^{\mathrm{som}}\right) = \mathbb{E}\left[\left|\frac{\partial\delta^C}{\partial x_k^{\mathrm{som}}}\right|\right] \mathrm{Cov}(x_k^{\mathrm{som}}, x_i^{\mathrm{som}}) \tag{41}$$

Therefore, if the neuron $i$ in question is the same neuron $k$ on which the state value estimation is based or if the neuron $i$ sufficiently covaries with the neuron $k$, the neuron $i$ is considered task-relevant. While it is challenging to write down a simple form of $\mathrm{Cov}(|\Delta\hat{V}|, x_i^{\mathrm{som}})$ for more complex models of $\hat{V}$, we can look at another case where $\hat{V}$ depends only on two neurons $k$ and $l$

$$\frac{\partial\delta^C}{\partial x_j^{\mathrm{som}}} = 0\,,\ \forall j \notin \{k, l\} \tag{42}$$

$$\mathrm{Cov}\left(|\Delta\hat{V}_2|, x_i^{\mathrm{som}}\right) = \mathrm{Cov}\left(\left|\frac{\partial\delta^C}{\partial x_k^{\mathrm{som}}} x_k^{som} + \frac{\partial\delta^C}{\partial x_l^{\mathrm{som}}} x_l^{som}\right|, x_i^{\mathrm{som}}\right) \tag{43}$$

Equation 43 tells us that, as long as the neurons $k$ and $l$ do not have reverse effects on $\hat{V}$ at the same time, the covariance can be considered to have distributive properties

$$\mathrm{Cov}\left(|\Delta\hat{V}_2|, x_i^{\mathrm{som}}\right) = \mathbb{E}\left[\left|\frac{\partial\delta^C}{\partial x_k^{\mathrm{som}}}\right|\right] \mathrm{Cov}(x_k^{\mathrm{som}}, x_i^{\mathrm{som}}) + \mathbb{E}\left[\left|\frac{\partial\delta^C}{\partial x_l^{\mathrm{som}}}\right|\right] \mathrm{Cov}(x_l^{\mathrm{som}}, x_i^{\mathrm{som}}) \tag{44}$$

However, if they do negate each other, their task relevance and the task relevance with which they covary, will be reduced. This is a desired effect since it prevents self-canceling activities in the state-value estimation model without any effective consequences on the final prediction $\hat{V}$ to be recognized as task relevant.

#### 3.3 Setting the plasticity threshold $\theta_i$

We can note that for conditioned salience to be effective, (i.e., apical weights increasing for the most task-relevant neurons and not for task-irrelevant neurons) $\rho_i^{th}$ should be in a certain range. At the very least, the most task-relevant neuron $k = \mathrm{argmax}_i(\rho_i)$ should become amplified and completely task-irrelevant neurons ($\rho_i \leq 0$) should not. Thus, we can write the broadest range of $\theta_i$ for the system to function properly as

$$0 < \rho_i^{th} = \frac{\theta_i}{\mathbb{E}[x_{i,t}^{\mathrm{som}}]} - 1 < \max_i \frac{\mathrm{Cov}([|\Delta \hat{V}_t|], x_{i,t}^{\mathrm{som}})}{\mathbb{E}[|\Delta \hat{V}_t|]\mathbb{E}[x_{i,t}^{\mathrm{som}}]} = \rho_i \quad (45)$$

This expression allows us to derive the range of acceptable values for $\theta_i$

$$\theta_i > \mathbb{E}[x_i^{\mathrm{som}}] \ , \ \forall i \quad (46)$$

$$\theta_i < \mathbb{E}_t[x_{i,t}^{\mathrm{som}}]\,(\rho_i + 1) \quad (47)$$

While in the pure-selectivity model a single global value for $\theta_i = 0.1$ for all $i$ was easy to find, large discrepancies in $\mathbb{E}_t[x_{i,t}^{\mathrm{som}}]$ for different neurons $i$ could render conditioned salience learning more challenging. For such cases, one could make the synaptic weight update rule in equation 16 sensitive to the expectation of the basal input by setting $\theta = \theta_0 \mathbb{E}[x_{i,t}^{\mathrm{som}}]$

$$\Delta w_{i,t}^{\mathrm{ap}} = \eta_w \ x_t^{\mathrm{pre}}\left(x_{i,t}^{\mathrm{som}} - \theta_0 \mathbb{E}\left[x_{i,t}^{\mathrm{som}}\right]\right) \quad (48)$$

In this case the constraints in equation 45 can be reformulated as

$$1 < \theta_0 < 1 + \frac{\mathrm{Cov}(\mathbb{E}_\pi[|\Delta \hat{V}_t|], x_{i,t}^{\mathrm{som}})}{\mathbb{E}_{t,\pi}[|\Delta \hat{V}_t|]\mathbb{E}_t[x_{i,t}]} = 1 + \rho_i \quad (49)$$

If $\theta_0 = 1$, all neurons will be amplified and if $\theta_0 > 1 + \rho_i$, none will be amplified. For the mixed-selectivity model, we selected $\theta_0 = 4$, which successfully amplified the most relevant neurons without amplifying the most irrelevant neurons (Supplementary Fig. 14c).
